# A disease-associated gene desert orchestrates macrophage inflammatory responses via ETS2

**DOI:** 10.1101/2023.05.05.539522

**Authors:** CT Stankey, C Bourges, T Turner-Stokes, AP Piedade, C Palmer-Jones, I Papa, M Dos Silva dos Santos, LO Randzavola, L Speidel, EC Parkes, W Edwards, AP Rochford, CD Murray, JI MacRae, P Skoglund, C Wallace, MZ Cader, DC Thomas, JC Lee

## Abstract

Increasing global rates of autoimmune and inflammatory disease present a burgeoning threat to human health^1^. This is compounded by the limited efficacy of available treatments^1^ and high failure rates during drug development^2^ – underscoring an urgent need to better understand disease mechanisms. Here we show how genetics could address this challenge. By investigating an intergenic haplotype on chr21q22, independently linked to inflammatory bowel disease (IBD), ankylosing spondylitis, primary sclerosing cholangitis and Takayasu’s arteritis^3–6^, we discover that the causal gene, *ETS2*, is a master regulator of inflammatory responses in human macrophages and delineate how the risk haplotype increases *ETS2* expression. Genes regulated by ETS2 were prominently expressed in affected tissues from chr21q22-associated diseases and more enriched for IBD GWAS hits than almost all previously described pathways. Overexpressing *ETS2* in resting macrophages produced an activated effector state that phenocopied intestinal macrophages from IBD^7^, with upregulation of multiple drug targets including TNFα and IL-23. Using a database of cellular signatures^8^, we identify drugs that could modulate this pathway and validate the potent anti-inflammatory activity of one class of small molecules *in vitro* and *ex vivo*. Together, this highlights the potential for common genetic associations to improve both the understanding and treatment of human disease.

---

Currently, nearly 5% of the world’s population are affected by at least one autoimmune or inflammatory disease. These heterogeneous conditions, which range from Crohn’s disease and ulcerative colitis (collectively IBD) to psoriasis and rheumatoid arthritis, share a common need for better treatments, but only ∼10% of drugs entering clinical development ever become approved therapies^2^. This high failure rate is principally due to a lack of efficacy^9^ or – put another way – because the pathways being targeted are less important than they were assumed to be. Genetics provides a unique opportunity to elucidate disease mechanisms, with hundreds of regions of the human genome now directly linked to the pathogenesis of one or more autoimmune or inflammatory disease^10^. Indeed, drugs that target pathways implicated by genetics have a substantially higher chance of becoming approved therapies^11^.

To fully realise the potential of genetics, however, knowledge of where risk variants lie must first be translated into an understanding of how they contribute to disease^10^. This is a formidable challenge since most disease-associated genetic variants do not lie in coding DNA, where effects on protein sequence/structure can be easily predicted, but in the enigmatic non-coding genome where the same DNA sequence can have different biological functions depending on the cell-type and/or external stimuli^10^. Most risk variants are thought to affect gene regulation^12^, but the need to identify the causal gene(s) – which may lie up to one million bases away – and the causal cell-type(s), which may only express the causal gene under specific conditions, have hindered attempts to discover disease mechanisms. For example, although genome-wide association studies (GWAS) have identified over 240 IBD risk loci^3^, fewer than 10 have been mechanistically resolved and, to date, none have led to new therapies.

## Molecular mechanisms at the pleiotropic chr21q22 locus

Several genetic variants predispose to more than one disease – highlighting both their biological importance and an opportunity to discover shared disease mechanisms. One notable example is an intergenic region on chr21q22, where the major (risk) allele haplotype has been independently associated with five different inflammatory diseases^3–6^. Although the associated locus does not contain any genes, there are several nearby candidates including *PSMG1*, *BRWD1* and *ETS2* (**Fig.1a**), all of which have been proposed to be potentially causal in previous studies^3–6, 13^. The underlying biological mechanisms, however, remain unknown. We hypothesised that this intergenic locus must contain a distal enhancer and – since the associated diseases are all immune-mediated despite affecting different organs – searched for evidence of enhancer activity in disease-relevant immune cell-types. Using H3K27ac ChIP-seq data, which marks active enhancers/promoters, we found that this locus contains a monocyte/macrophage-specific enhancer (**Fig.1a**). Monocytes and monocyte-derived macrophages play a central role in the pathogenesis of many autoimmune and inflammatory diseases, producing cytokines that are often targeted by the most effective therapies^14^.

**Figure 1.**
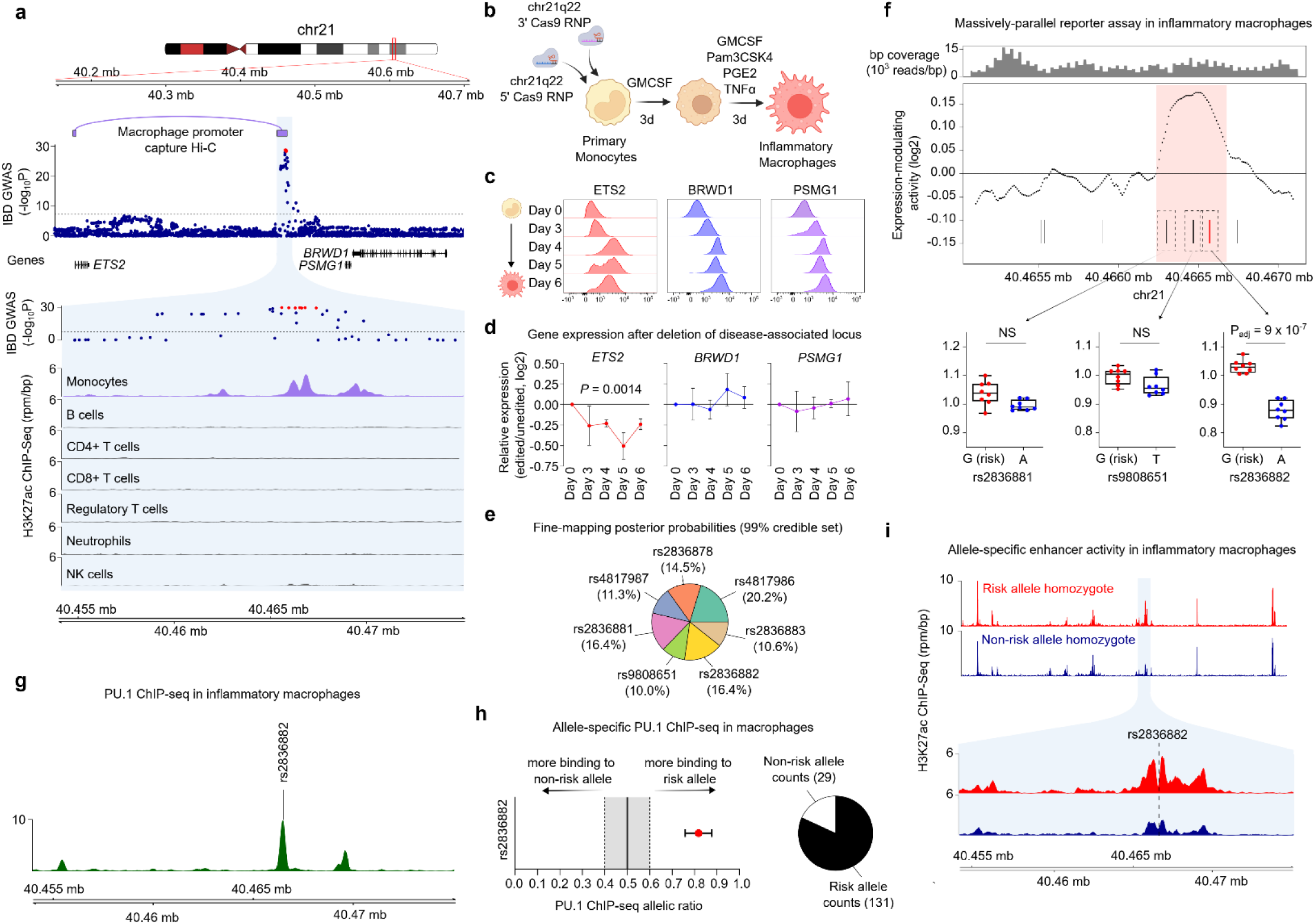
Resolving molecular mechanisms at chr21q22. **a**. Annotation of the disease-associated chr21q22 locus depicting the IBD genetic association, physical interactions of the disease-associated haplotype in macrophages (promoter-capture Hi-C), and H3K27ac ChIP-seq data from various immune cell-types. **b.** Schematic of experiment to determine the function of the chr21q22 locus in monocyte-derived macrophages polarised under chronic inflammatory (“TPP”) conditions. **c.** Histograms depicting the expression of *ETS2*, *BRWD1*, and *PSMG1* during inflammatory macrophage polarisation, measured using PrimeFlow RNA assays that quantify RNA by flow cytometry. Data are representative of one of four donors. **d.** Relative *ETS2*, *BRWD1*, and *PSMG1* expression in chr21q22-edited inflammatory macrophages (relative to non-targeting control cells; NTC). Plot shows log2 fold-change in mean fluorescence intensity (n=4, data represent mean±SEM, two-way ANOVA). **e.** SuSiE fine-mapping posterior probabilities for IBD-associated SNPs at the 21q22 locus (99% credible set). **f.** MPRA at the chr21q22 locus depicting oligonucleotide coverage (top), enhancer activity in inflammatory macrophages (analysed using a sliding window analysis of tiling oligos; middle), and expression-modulating effects of candidate SNPs within the identified enhancer (bottom) (n=8). Shaded region in enhancer activity plot indicates region of significant enhancer activity. **g.** PU.1 ChIP-seq peaks at the chr21q22 locus in macrophages. **h.** BaalChIP analysis of allele-specific PU.1 binding at rs2836882 in two heterozygous macrophage datasets (data represent 95% posterior distribution of allelic binding ratio) **i.** H3K27ac ChIP-seq data from major (top) or minor (bottom) allele homozygotes at the chr21q22 locus. Data are representative of two of four donors.

To identify the gene regulated by this enhancer, we first considered publicly-available data from human monocytes, including promoter-capture Hi-C^15^ and eQTL datasets^16^. We found that the disease-associated locus physically interacts with the promoter of *ETS2*, the most distant of the candidate genes (located 290-kb away) and that the risk haplotype correlates with higher *ETS2* expression, especially after monocyte activation (**Extended Data Fig.1**). Interestingly, however, the disease association was predicted to share a causal variant not with the strong eQTL in activated monocytes, but rather with a weaker eQTL in resting monocytes (**Extended Data Fig.1**). To directly confirm the identity of the target gene, we deleted the 1.85-kb enhancer region in primary human monocytes using CRISPR-Cas9 ribonucleoprotein (RNP) complexes containing gRNAs that flank the enhancer (**Fig.1b**, **Extended Data Fig.2**). These cells were then cultured with combination of inflammatory mediators, including TNFα (a pro-inflammatory cytokine), prostaglandin E2 (a pro-inflammatory lipid) and Pam3CSK4 (a TLR1/2 agonist). This model, termed “TPP”, was designed to mimic a chronic inflammatory environment^17^, and better recapitulates the state of patient-derived monocytes/macrophages than classical M1 or M2 models (ref.18 and **Extended Data Fig.2**). Because flow cytometry antibodies were not available for any of the candidate genes, we used PrimeFlow to measure the dynamics of RNA transcription and found that expression of all 3 genes (*ETS2*, *BRWD1*, *PSMG1*) increased in unedited cells upon exposure to inflammatory stimuli (**Fig.1c**). Deletion of the chr21q22 enhancer did not affect the increase in *BRWD1* and *PSMG1* expression, but the upregulation of *ETS2* was significantly reduced (**Fig.1d**) – confirming that this pleiotropic locus functions as a distal *ETS2* enhancer in monocytes and monocyte-derived macrophages.

We next sought to discover the variant responsible for disease risk at the chr21q22 locus. Unfortunately, statistical fine-mapping – using the largest IBD GWAS to date^3^ – could not identify the causal variant due to high linkage disequilibrium between the candidate single nucleotide polymorphisms (SNPs) (**Methods**, **Fig.1e**). We therefore used a high-throughput functional approach to first delineate the active enhancers at the locus, and then determine if any candidate SNPs within these regulatory regions might alter enhancer activity. This method – massively-parallel reporter assay (MPRA) – can simultaneously characterise enhancer activity in thousands of short DNA sequences by coupling each to a uniquely-barcoded reporter gene within an expression vector^19^. Genetic sequences that modulate gene expression can be identified by normalising the barcode counts in mRNA extracted from transfected cells to their equivalent counts in the input DNA library. After adapting the MPRA vector for use in primary macrophages (**Methods**, **Extended Data Fig.3**), we synthesised a pool of overlapping oligonucleotides (oligos) to tile the 2-kb region encompassing all candidate SNPs at chr21q22, and included additional oligos containing either risk or non-risk alleles for every variant. The resulting vector library was transfected into inflammatory macrophages from multiple donors – thus ensuring that a physiological repertoire of transcription factors would be present to interact with the chr21q22 genomic sequences. Using a sliding window analysis to map active enhancers across the tiling sequences, we identified a single 442-bp region of enhancer activity (chr21:40466236-40466677, hg19; **Fig.1f**) that harboured three (of seven) candidate SNPs. Two of these polymorphisms were transcriptionally inert, but the third (rs2836882) had the strongest expression-modulating effect of any candidate variant, with the risk allele (G) significantly increasing transcription – consistent with the known eQTL (**Fig.1f**). Examining rs2836882 further, we noticed that this SNP lay within an experimentally-confirmed PU.1 ChIP-seq peak in human macrophages (**Fig.1g**). PU.1 is an important myeloid pioneer factor^20^ that can bind to heterochromatin, initiate nucleosome remodelling – thus enabling other transcription factors to bind – and activate transcription^21^. To determine whether rs2836882 might affect PU.1 binding, we identified two publicly-available macrophage PU.1 ChIP-seq datasets from rs2836882 heterozygotes and used BaalChIP^22^ to assess for allelic imbalances in PU.1 binding. Despite not lying within a canonical PU.1 binding motif, significant allele-specific PU.1 binding was detected at rs2836882, with over 4-fold greater binding to the risk allele in both datasets (**Fig.1h**). This result was replicated in TPP macrophages from five heterozygous donors by immunoprecipitating PU.1 and genotyping the bound DNA (**Extended Data Fig.4**). Together, this suggests that the rs2836882 risk allele should increase enhancer activity, consistent with the MPRA and eQTL results.

To test whether allele-specific enhancer activity was evident at the endogenous locus, we performed H3K27ac ChIP-seq in inflammatory macrophages from two rs2836882 major allele homozygotes and two minor allele homozygotes. While several nearby enhancer peaks were similar between these donors, the enhancer activity overlying rs2836882 was considerably stronger in major (risk) allele homozygotes (**Fig.1i**), contributing to a ∼2.5-fold increase in enhancer activity across the extended chr21q22 locus (**Extended Data Fig.4**). Collectively, these data reveal a genetic mechanism whereby the putative causal variant at chr21q22 – identified via its functional consequences in primary macrophages – promotes binding of a pioneer transcription factor and increases the activity of a long-range *ETS2* enhancer.

## ETS2 is essential for macrophage inflammatory responses

Having identified a plausible mechanism by which the chr21q22 risk haplotype increases *ETS2* expression, we sought to better understand the role of *ETS2* in monocytes/macrophages. *ETS2* is a member of the ETS family of transcription factors, which has been mainly studied as a proto-oncogene in cancer^23^. In contrast, the role of *ETS2* in primary human macrophages has been less clearly defined, with previous studies using either cell-lines or complex mouse models and largely focusing on a limited number of downstream molecules^24–28^. This has led to contradictory reports, with *ETS2* being described as both necessary and redundant for macrophage development^29, 30^, and both pro- and anti-inflammatory^24–28^. To elucidate the specific role of *ETS2* in inflammatory human macrophages – and determine how dysregulated *ETS2* expression might contribute to disease – we first used a CRISPR-Cas9-based loss-of-function approach (**Fig.2a**). Two gRNAs targeting different *ETS2* exons were designed, validated and individually incorporated into Cas9 RNPs for transfection into primary monocytes – thereby minimising the chance that any effect was due to off-target editing. These gRNAs resulted in on-target editing in ∼89% and ∼79% of total cells respectively (**Extended Data Fig.2**). No differences in cell viability or expression of macrophage markers were observed with either gRNA, suggesting that *ETS2* was not required for inflammatory macrophage survival or differentiation (**Extended Data Fig.2**). In contrast, production of pro-inflammatory cytokines, including IL-6, IL-8 and IL-1β, was significantly reduced following *ETS2* disruption (**Fig.2b**), whereas IL-10 – an anti-inflammatory cytokine – was less affected. TNFα secretion could not be assessed as it was included in the differentiation culture. We therefore investigated whether *ETS2* was also required for other macrophage effector functions. First, we examined phagocytosis using fluorescently-labelled particles that can be detected by flow cytometry. Similar to pro-inflammatory cytokine production, phagocytosis was significantly impaired following *ETS2* disruption (**Fig.2c**). We next measured extracellular reactive oxygen species (ROS) production – a key effector response that directly contributes to tissue damage in inflammatory disease^31^. Disrupting *ETS2* profoundly reduced the oxidative burst following macrophage activation – an effect that appeared to be due to reduced expression of key NADPH oxidase components (**Fig.2d**, **Extended Data Fig.5**). Together, this suggests that *ETS2* is required for multiple effector functions in inflammatory macrophages.

**Figure 2.**
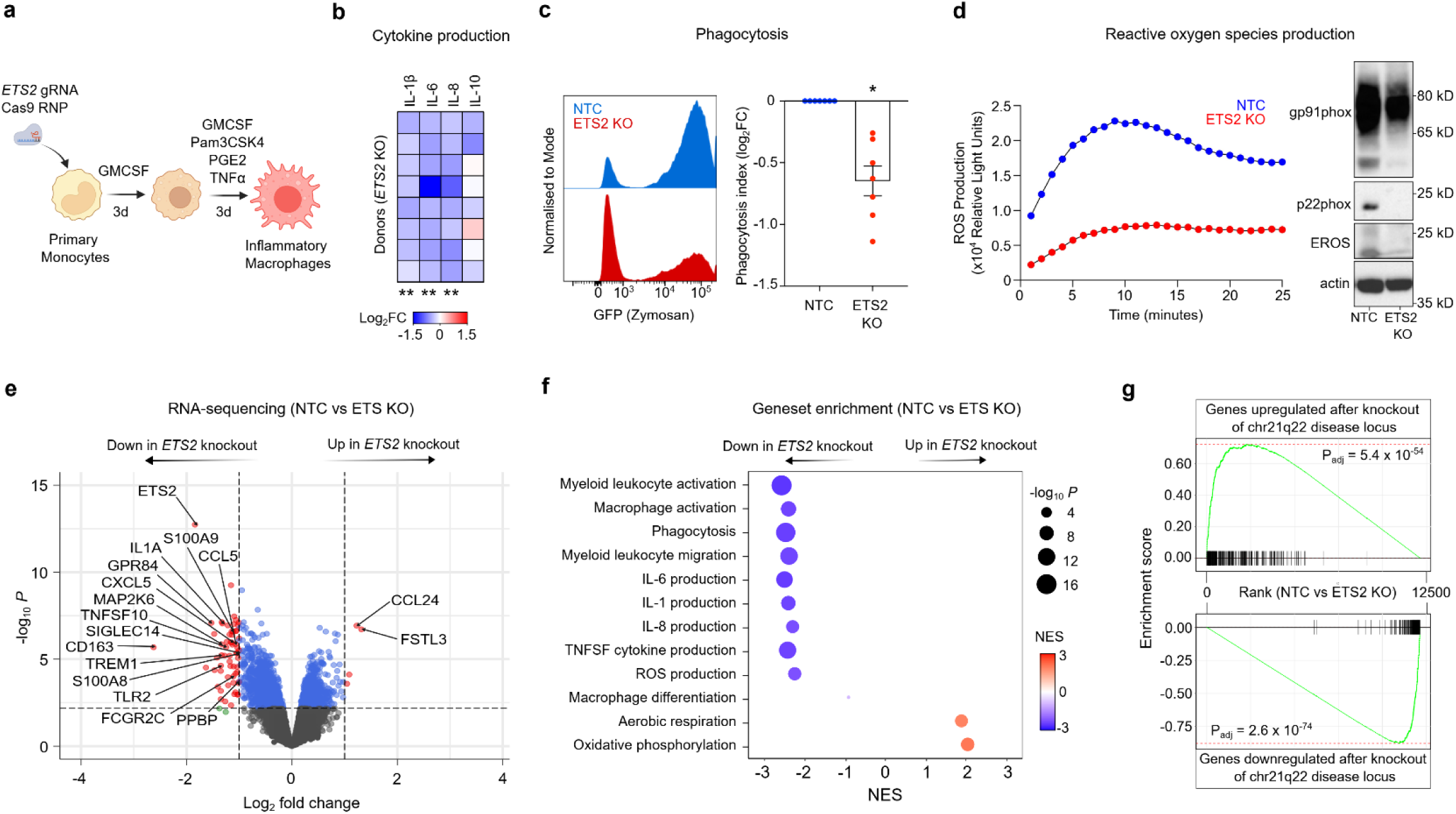
*ETS2* is essential for macrophage inflammatory responses. **a.** Schematic of experiment for disrupting *ETS2* in primary monocytes and differentiating monocyte-derived macrophages under chronic inflammatory conditions. **b.** Macrophage cytokine secretion following *ETS2* disruption. Heatmap shows log2 fold-change of cytokine concentrations in the supernatants of *ETS2*-edited macrophages relative to unedited macrophages transfected with a non-targeting control gRNA-containing RNP (NTC). n=9, Wilcoxon matched-pairs test, two-tailed. **c.** Histogram depicting phagocytosis of fluorescently-labelled zymosan particles by *ETS2*-edited and unedited macrophages (left). Data representative of one of seven donors. Phagocytosis index in *ETS2*-edited and unedited macrophages (calculated as product of proportion and mean fluorescence intensity of phagocytosing cells; right). Plot depicts log2 fold-change in phagocytosis index for *ETS2*-edited macrophages relative to unedited cells (Wilcoxon signed-rank test, two-tailed; data represent mean±SEM). **d.** Production of ROS by *ETS2*-edited and unedited inflammatory macrophages (measured in relative light units; left). Data representative of one of six donors. Western blot for gp91*phox*, p22*phox*, and EROS expression in *ETS2*-edited and unedited macrophages (right). Data representative of one of three donors. **e.** Differentially-expressed genes in *ETS2*-edited versus unedited inflammatory macrophages (limma with voom transformation, n=8). **f.** Gene set enrichment analysis (fGSEA) of differentially-expressed genes between *ETS2*-edited and unedited inflammatory macrophages. Results of selected Gene Ontology Biological Pathways shown. Dot size represents *P*-value and colour denotes normalised enrichment score (NES). **g.** Enrichment of differentially-expressed genes following deletion of the disease-associated chr21q22 locus (upregulated genes, top; downregulated genes, bottom) in *ETS2*-edited versus unedited macrophages. * *P* < 0.05, ** *P* < 0.01.

To better understand the molecular basis for these distinct functional effects, we performed RNA-sequencing (RNA-seq) in *ETS2*-edited and unedited inflammatory macrophages from multiple donors. Disruption of *ETS2* led to widespread transcriptional changes, with reduced expression of many inflammatory genes, including several well-known initiators and amplifiers of inflammation (**Fig.2e**). Affected gene classes included cytokines (e.g. *TNFSF10/TRAIL*, *TNFSF13*, *IL1A, IL1B*), chemokines (e.g. *CXCL1*, *CXCL3*, *CXCL5*, *CCL2, CCL5*), secreted effector molecules (e.g. *S100A8*, *S100A9*, *MMP14*, *MMP9*), cell surface receptors (e.g. *FCGR2A*, *FCGR2C, TREM1*), pattern recognition receptors (e.g. *TLR2*, *TLR6*, *NOD2*), and signalling molecules (e.g. *MAP2K*, *GPR84, NLRP3*). To better characterise the pathways affected by *ETS2* deletion, we performed gene-set enrichment analysis (fGSEA) using the Gene Ontology Biological Pathways dataset. This corroborated the observed functional effects (**Fig.2f**), with the most negatively-enriched pathways (downregulated following *ETS2* disruption) relating to macrophage activation, pro-inflammatory cytokine production, phagocytosis and ROS production. Genes involved in macrophage migration were also significantly downregulated, but gene sets relating to monocyte-to-macrophage differentiation were not significantly affected – consistent with *ETS2* being required for inflammatory effector functions, but not influencing monocyte-to-macrophage development. Although fewer genes increased in expression following *ETS2* deletion (**Fig.2e**), positive enrichment was noted for genes involved in aerobic respiration and oxidative phosphorylation (OXPHOS; **Fig.2f**) – metabolic processes linked to anti-inflammatory macrophage behaviour^32^. Collectively, these data identify an indispensable role for *ETS2* in a range of macrophage effector functions, which could explain why dysregulated *ETS2* expression is associated with multiple inflammatory diseases. Indeed, deletion of the disease-associated chr21q22 enhancer phenocopied both the transcriptional and functional consequences of disrupting *ETS2* (**Fig.2g**, **Extended Data Fig.5**).

## ETS2 orchestrates macrophage inflammatory responses

Having found that *ETS2* was essential for monocyte-derived macrophage effector functions, we next investigated whether it might also be sufficient to drive them – as would be expected of a master regulator of inflammatory responses. This was particularly important because although loss-of-function approaches can identify a gene’s biological role(s), the chr21q22 risk haplotype increases *ETS2* expression. To do this, we first optimised a method for controlled overexpression of target genes in primary macrophages by transfecting defined amounts of *in vitro* transcribed mRNA that was modified to minimise immunogenicity (**Fig.3a**, **Extended Data Fig.3, Methods**). Resting, non-activated (M0) macrophages were then transfected with mRNA encoding *ETS2* or its reverse complement – thereby controlling for quantity, length and purine/pyrimidine composition of the transfected mRNA but with a transcript that would not be translated (**Fig.3b**). After transfection, cells were exposed to low-dose lipopolysaccharide for 6 hours to initiate a low-grade inflammatory response that could be amplified if *ETS2* was sufficient to drive inflammatory responses (**Fig.3a**). We first quantified secreted cytokines and found that *ETS2* overexpression increased production of several pro-inflammatory cytokines, although IL-10 was again less affected (**Fig.3c**). To better characterise the consequences of *ETS2* overexpression, we performed RNA-seq and examined the macrophage activation pathways that had required *ETS2*. Strikingly, all of these inflammatory pathways – including macrophage activation, pro-inflammatory cytokine production, ROS production, phagocytosis and migration – were induced in a dose-dependent manner following *ETS2* overexpression, with greater enrichment of every pathway when more *ETS2* mRNA was transfected (**Fig.3d**). This shows that *ETS2* is both necessary and sufficient for inflammatory responses in human macrophages, consistent with being a master regulator of effector functions during chronic inflammation, whose dysregulation is directly linked to human disease.

**Figure 3.**
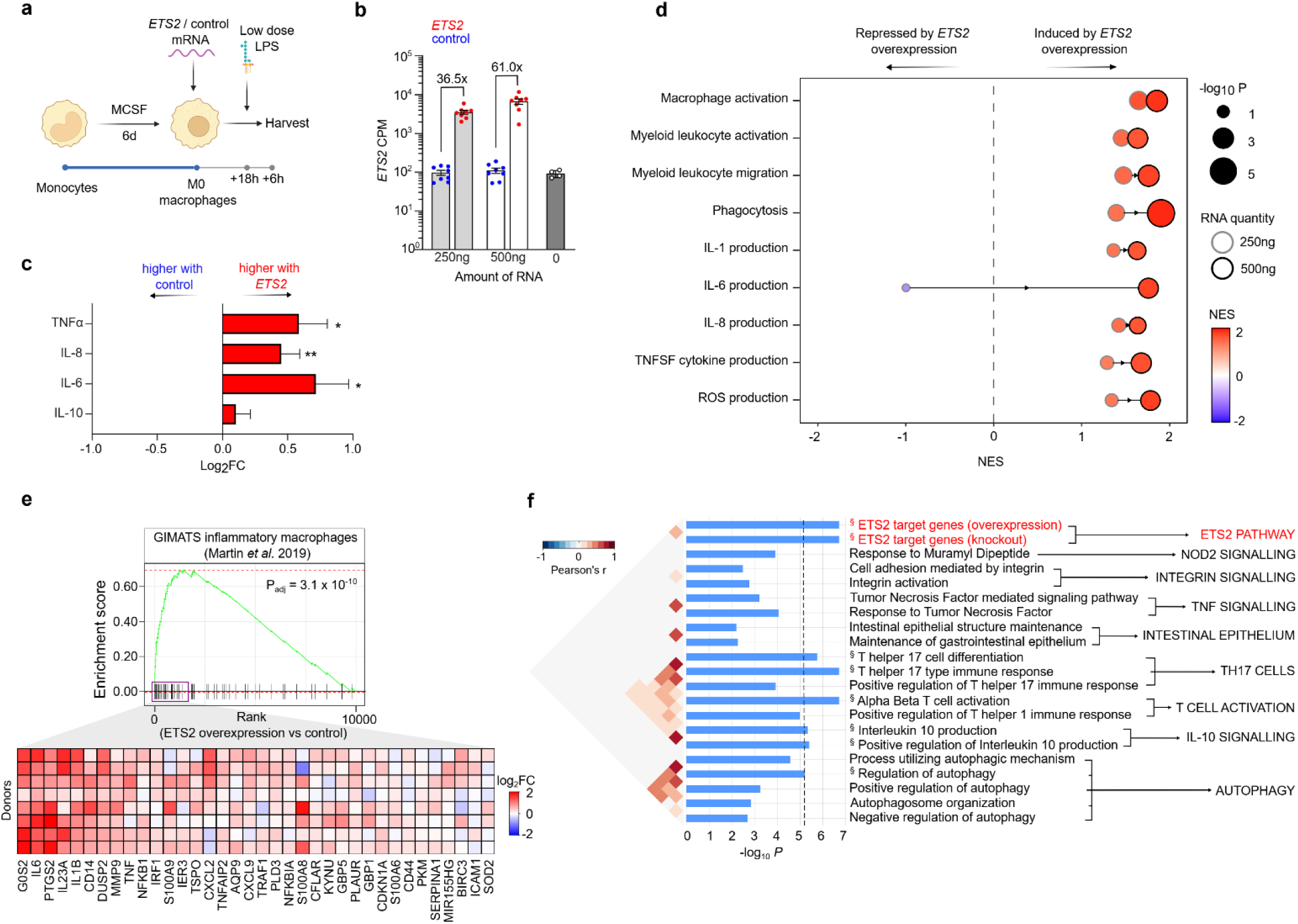
ETS2 orchestrates macrophage inflammatory responses. **a.** Schematic of *ETS2* overexpression experiment. Resting (M0) human macrophages were transfected with pre-defined amounts of *in vitro* transcribed *ETS2* mRNA or control mRNA (*ETS2* reverse complement), activated with low dose LPS (0.5ng/ml), and harvested. n=8. **b.** *ETS2* mRNA levels in macrophages transfected with *ETS2* or control mRNA or untransfected (from separate experiment). **c.** Cytokine secretion following *ETS2* overexpression. Plot shows log2 fold-change of cytokine concentrations in macrophage supernatants (*ETS2* relative to control) following transfection with 500ng mRNA. **d.** Gene set enrichment analysis (fGSEA) of differentially-expressed genes between *ETS2*-overexpressing and control macrophages. Results shown for the same Gene Ontology Biological Pathways that were negatively enriched following ETS2 editing. Dot size represents *P*-value, colour denotes normalised enrichment score (NES), and border colour denotes amount of transfected mRNA. **e.** Enrichment of a disease-associated inflammatory macrophage gene signature, derived from single cell RNA-seq of Crohn’s disease intestinal biopsies, in *ETS2*-overexpressing macrophages (relative to control; top). Heatmap of leading-edge genes showing log2 fold-change of gene expression in *ETS2*-overexpressing macrophages relative to control (500ng mRNA; bottom). **f.** SNPsea analysis of enrichment of 241 IBD-associated loci within *ETS2*-regulated genes (red) and pathways previously linked to IBD pathogenesis (black). Significantly enriched pathways (Bonferroni-corrected permutation *P* < 0.05) indicated by §. * *P* < 0.05, ** *P* < 0.01.

## ETS2-regulated genes play a central role in IBD

To understand whether *ETS2* might contribute to the macrophage phenotype observed in inflammatory disease, we compared the transcriptional consequences of overexpressing *ETS2* with a gene signature of intestinal macrophages from Crohn’s disease – one of the conditions associated with the chr21q22 locus. Single cell RNA-seq analysis has previously shown that active Crohn’s disease is characterised by an expanded population of inflammatory monocyte-derived macrophages that contributes to anti-TNFα resistance^7^. Using the Crohn’s disease macrophage signature as a gene set, we found that overexpressing *ETS2* in resting macrophages induced a transcriptional state that closely resembled disease macrophages, with core (“leading edge”) enrichment of the majority of genes in the signature, including many that encode therapeutic targets (**Fig.3e**).

Given the importance of *ETS2* in macrophage responses, and the fact that *ETS2* overexpression phenocopied a disease-associated inflammatory state, we hypothesised that other genetic associations might also implicate this previously uncharacterised pathway. An important goal of GWAS was to identify key disease pathways^10^, but this has proven challenging due to a paucity of confidently-identified causal genes and a limited understanding of how these are affected by genetic variation^10^. To better characterise the genetic risk attributable to the macrophage ETS2 pathway, we focused on IBD since this has far more genetic associations than any other chr21q22-associated disease. Examining the list of commonly downregulated genes following *ETS2* editing (P_adj_ < 0.05 for both gRNAs), we identified over 20 IBD risk genes – including many that have been proposed to be causal at their respective loci^3, 33^ (**Extended Data Table 1**). These included genes that are thought to affect macrophage biology (e.g. *SP140*, *LACC1/FAMIN*, *CCL2*, *CARD9*, *CXCL5, TLR4, SLAMF8, FCGR2A*) as well as some that are highly expressed in macrophages but not previously linked to specific pathways (e.g. *ADCY7*, *PTPRC*, *TAGAP*, *PTAFR*, *PDLIM5*). To more formally assess the enrichment of an ETS2-regulated inflammatory pathway in IBD genetics – and compare this to known disease pathways – we used SNPsea^34^, an algorithm designed to identify pathways affected by disease loci. 241 IBD-associated genetic loci were tested for enrichment in 7,660 pathways, comprising 7,658 Gene Ontology Biological Pathways and 2 overlapping lists of ETS2-regulated genes (either those downregulated following *ETS2* editing or upregulated following *ETS2* overexpression). Significance of enrichment was empirically computed using 5 million matched null SNP sets, and disease pathways previously implicated by genetics were extracted for comparison. Strikingly, ETS2 target genes – however they were defined – were more strongly enriched for IBD-associated loci than almost all previously implicated pathways, with not a single null SNP set showing greater enrichment in either of the ETS2-regulated gene lists. After applying a stringent Bonferroni multiple-testing correction, only ETS2-regulated genes and IBD pathways relating to T cell activation, T-helper 17 cells, autophagy and IL-10 signalling showed significant enrichment (**Fig.3f**). This suggests that ETS2 signalling in macrophages plays a fundamental role in IBD pathogenesis, with stronger genetic enrichment than most previously implicated pathways.

## ETS2 controls inflammatory responses via transcriptional and metabolic effects

We next sought to understand how *ETS2* controlled such diverse macrophage effector functions. Studying ETS2 biology is challenging because no ChIP-seq-grade antibodies exist, precluding direct identification of its transcriptional targets. Even the ENCODE project, which performed ChIP-seq for 181 transcription factors, was unable to directly immunoprecipitate ETS2^35^. We therefore first used a “guilt-by-association” approach to identify genes that were co-expressed with *ETS2* across 64 different human macrophage polarisation conditions^17^. This identified *PFKFB3* – encoding the rate-limiting enzyme of glycolysis – as the most highly co-expressed gene, with *HIF1A* also highly co-expressed (**Fig.4a**). Together, these genes are known to facilitate a “glycolytic switch” that is required for myeloid inflammatory responses^36^. We therefore hypothesised that *ETS2* might control inflammatory responses via metabolic reprogramming – a possibility supported by OXPHOS genes being negatively correlated with *ETS2* expression (**Fig.4a**) and upregulated following *ETS2* disruption (**Fig.2f**). To assess the metabolic consequences of disrupting *ETS2*, we quantified label incorporation from ^13^C-glucose in edited and unedited inflammatory macrophages using gas chromatography–mass spectrometry (GC-MS). Widespread modest reductions in labelled and total glucose metabolites were detected following *ETS2* disruption (**Fig.4b, Extended Data Fig.6**). This affected both glycolytic and TCA cycle metabolites, with significant reductions in intracellular and secreted lactate, a hallmark of anaerobic glycolysis, and succinate, an important inflammatory signalling metabolite^37^. These results would be consistent with glycolytic suppression, with reductions in TCA metabolites being due to reduced flux into TCA and increased consumption by mitochondrial OXPHOS^38^. To determine whether metabolic changes were responsible for ETS2-mediated inflammatory effects, we treated *ETS2*-edited macrophages with roxadustat, a HIF1α stabiliser that promotes glycolysis via HIF1α-mediated metabolic reprogramming. This had the predicted effect on genes involved in glycolysis and OXPHOS, but did not rescue the effects of *ETS2* disruption, either transcriptionally or functionally (**Fig.4c, Extended Data Fig.6**). Thus, while disrupting *ETS2* does alter glycometabolism, this does not fully explain the observed differences in inflammation.

**Figure 4.**
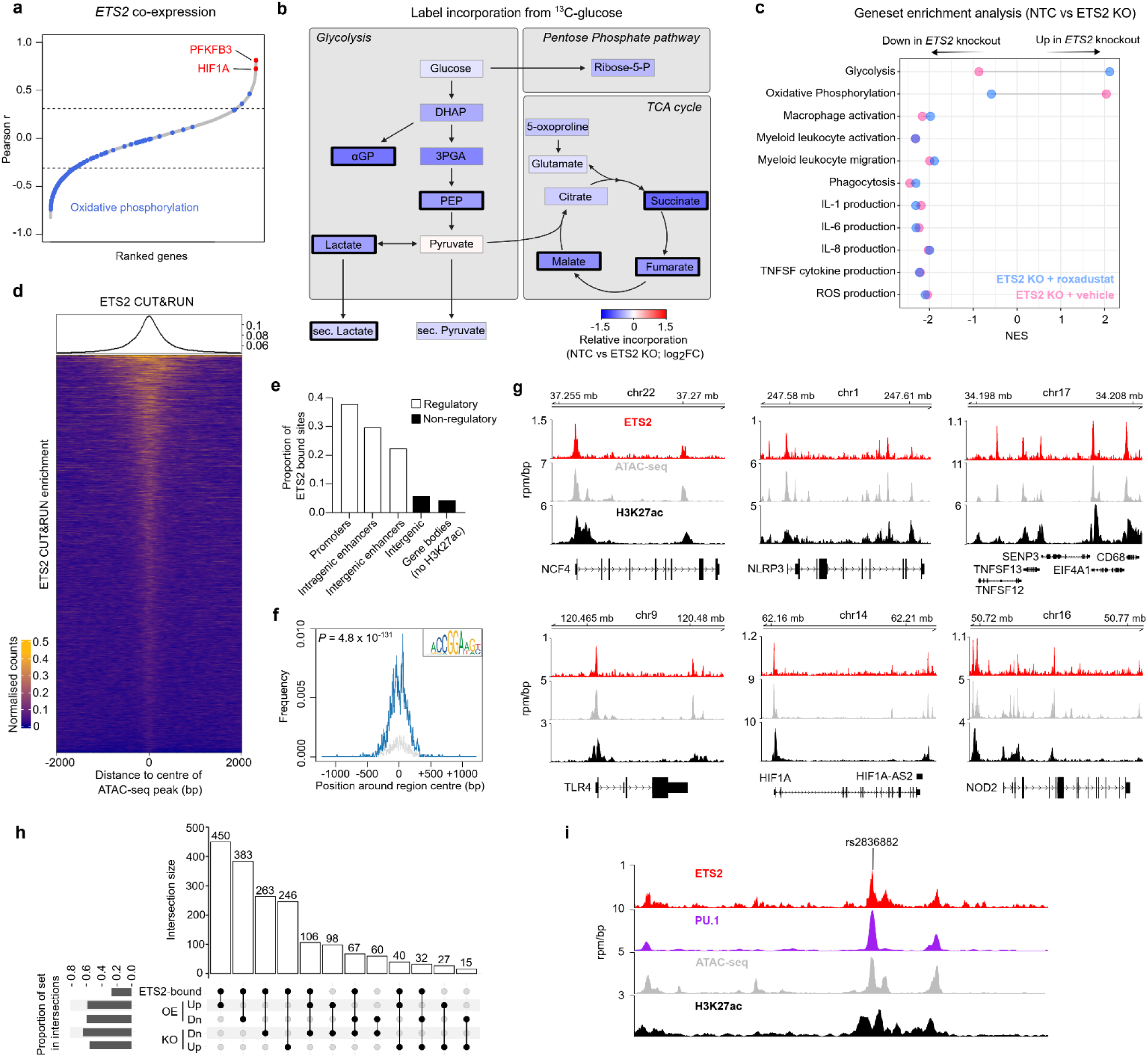
ETS2 directs macrophage responses via transcriptional and metabolic effects. **a.** Genes co-expressed with *ETS2* in 64 monocyte-derived macrophage datasets. Dotted lines equivalent to FDR *P* < 0.05. **b.** Effect of *ETS2* disruption on glucose metabolism. Colour denotes median log2 fold-change in label incorporation from ^13^C-glucose in *ETS2*-edited cells relative to unedited cells. Bold black border denotes *P* < 0.05 (Wilcoxon matched-pairs, two-tailed, n=6). **c.** Gene set enrichment analysis (fGSEA) of differentially-expressed genes between *ETS2*-edited and unedited TPP macrophages, treated with either roxadustat or vehicle. Results for selected Gene Ontology Biological Pathways shown. **d.** Enrichment heatmap of *ETS2* CUT&RUN peaks (IDR cut-off 0.01, n=2) in accessible chromatin from TPP macrophages (4-kb regions centred on ATAC-seq peaks). **e.** Features of ETS2 binding sites (based on gene coordinates and H3K27ac ChIP-seq in TPP macrophages). **f.** Enrichment of an ETS2 binding motif in ETS2 CUT&RUN peaks (hypergeometric P-value). **g.** ETS2 binding, chromatin accessibility (ATAC-seq), and enhancer activity (H3K27ac) at selected loci. **h.** UpSet plot of intersections between ETS2 gene lists, including genes with ETS2 peaks in their core promoters or cis-regulatory elements and significantly up-(Up) or down-regulated (Dn) genes following ETS2 editing (KO) or overexpression (OE). Vertical bars denote shared genes between lists, indicated by connected dots in lower panel. Horizontal bars denote proportion of gene list within intersections. **i.** ETS2 binding, chromatin accessibility (ATAC-seq), and enhancer activity (H3K27ac) at the disease-associated chr21q22 locus.

To try to elucidate the mechanism by which ETS2 controlled such diverse inflammatory effects, we revisited whether we could directly identify ETS2 target genes. Using a range of anti-ETS2 antibodies, we confirmed that none worked for ChIP-seq (data not shown) so investigated whether any might work for Cleavage-Under-Targets-and-Release-Using-Nuclease (CUT&RUN), which does not require formaldehyde fixation. One antibody identified multiple, significantly-enriched genomic regions (peaks) of which 6,560 were reproducibly detected across two biological replicates (Irreproducible Discovery Rate cut-off 0.01) with acceptable quality metrics^39^ (**Fig.4d**). These peaks were mostly located in active regulatory regions (90% in promoters or active enhancers, **Fig.4e**) and were highly enriched for a canonical ETS2 binding motif (4.02-fold enrichment over global controls, **Fig.4f**) – consistent with being sites of ETS2 binding. After combining the biological replicates to improve peak detection, we identified ETS2 binding at the promoters of many inflammatory genes, including several that are essential for distinct macrophage functions, such as *NCF4* (encoding a key NADPH oxidase component), *NLRP3* (encoding a key inflammasome component), and *TLR4* (encoding a key pattern recognition receptor) (**Fig.4g**). Overall, 48.3% of genes dysregulated following *ETS2* disruption, and 50.3% of genes dysregulated following *ETS2* overexpression, contained an ETS2 binding peak within their core promoter or putative cis-regulatory elements (**Fig.4h**) – consistent with ETS2 directly regulating a range of macrophage inflammatory responses. Notably, these gene targets included *HIF1A*, *PFKFB3*, and other glycolytic genes (e.g. *GPI*, *HK2*, and *HK3*), suggesting that the observed metabolic changes might be directly induced by ETS2, rather than being solely attributable to differences in inflammation. Intriguingly, we also detected ETS2 binding at its own enhancer at chr21q22 (**Fig.4i**). This is consistent with reports that PU.1 and ETS2 can interact synergistically^40^, and would implicate a feed-forward mechanism at the disease-associated locus, where increased *ETS2* expression reinforces *ETS2* enhancer activity. Together, these data implicate ETS2 as a master regulator of monocyte/macrophage responses during chronic inflammation, capable of directing a multifaceted effector programme, and creating a metabolic environment that is permissive for inflammation.

## ETS2-driven inflammation is evident in diseased tissue and can be targeted pharmacologically

The strong enrichment of IBD GWAS hits within ETS2-regulated genes led us to hypothesise that the transcriptional footprint of this pathway might be generally detectable in the affected tissues of chr21q22-associated diseases – a possibility that would have important therapeutic implications. Using publicly-available gene expression data from diseases linked to chr21q22 – intestinal macrophages from IBD, synovium from ankylosing spondylitis (AS), and liver from PSC – we confirmed that diseased tissue was significantly enriched for ETS2-regulated genes (**Fig.5a, Extended Data Fig.7**). We therefore investigated whether this pathway could be pharmacologically targeted. Specific ETS2 inhibitors do not exist and structural analyses indicate that there is no allosteric inhibitory mechanism that could be easily targeted^41^. We therefore used the NIH LINCS database to identify drugs that might modulate ETS2 activity^8^. This repository contains over 30,000 differentially-expressed genelists from cell-lines exposed to over 6,000 small molecules. Using fGSEA, 906 drug signatures were found to mimic the transcriptional effect of disrupting *ETS2* in inflammatory macrophages (P_adj_ < 0.05), including several approved treatments for IBD and AS (e.g. JAK inhibitors). Of these candidate therapies, the most common class were MEK inhibitors (**Fig.5b**), which are already licensed for non-inflammatory human diseases (e.g. neurofibromatosis). This result was not due to a single compound, but rather a class effect with multiple MEK1/2 inhibitors downregulating ETS2 target genes (**Fig.5c**). This made biological sense, since MEK1 and MEK2 – together with several other targets identified – are known upstream regulators of ETS-family transcription factors (**Fig.5d**). Indeed, some of these drug classes, including MEK1/2 and HSP90 inhibitors, have been reported to be beneficial in animal colitis models, although this is often a poor indicator of clinical efficacy – with several approved IBD treatments being ineffective in mice and many drugs that improve mouse models being ineffective in human IBD^42^. To determine whether MEK inhibition would abrogate ETS2-driven inflammatory responses in primary human macrophages, we differentiated monocytes under chronic inflammatory conditions and treated them with a selective, non-ATP competitive MEK inhibitor (PD-0325901; **Fig.5e**). We observed potent anti-inflammatory activity that phenocopied the effect of disrupting *ETS2* or deleting the chr21q22 enhancer (**Fig.5f, Extended Data Fig.8**), with downregulation of multiple ETS2-regulated pathways (including several drug targets; **Fig.5g**). To further explore the therapeutic potential of targeting ETS2 signalling^42^, we employed a human gut explant model. Intestinal mucosal biopsies were obtained from patients with active IBD, who were not receiving immunosuppressive or biologic therapies, and cultured with either a MEK inhibitor or a negative or positive control (**Methods**). Release of multiple IBD-associated inflammatory cytokines was significantly reduced by MEK inhibition – to comparable levels observed with infliximab (an anti-TNFα antibody widely used for IBD; **Fig.5h**). Moreover, we confirmed that expression of ETS2-regulated genes was reduced (**Fig.5i, Extended Data Fig.8)** and that there was significant improvement in a validated transcriptional inflammation score^43^ that reflects IBD-associated inflammation and has been shown to reduce upon effective therapy (**Fig.5j**). Together, this shows that targeting an upstream regulator of *ETS2* can abrogate pathological inflammation in a chr21q22-associated disease, and may be useful therapeutically.

**Figure 5.**
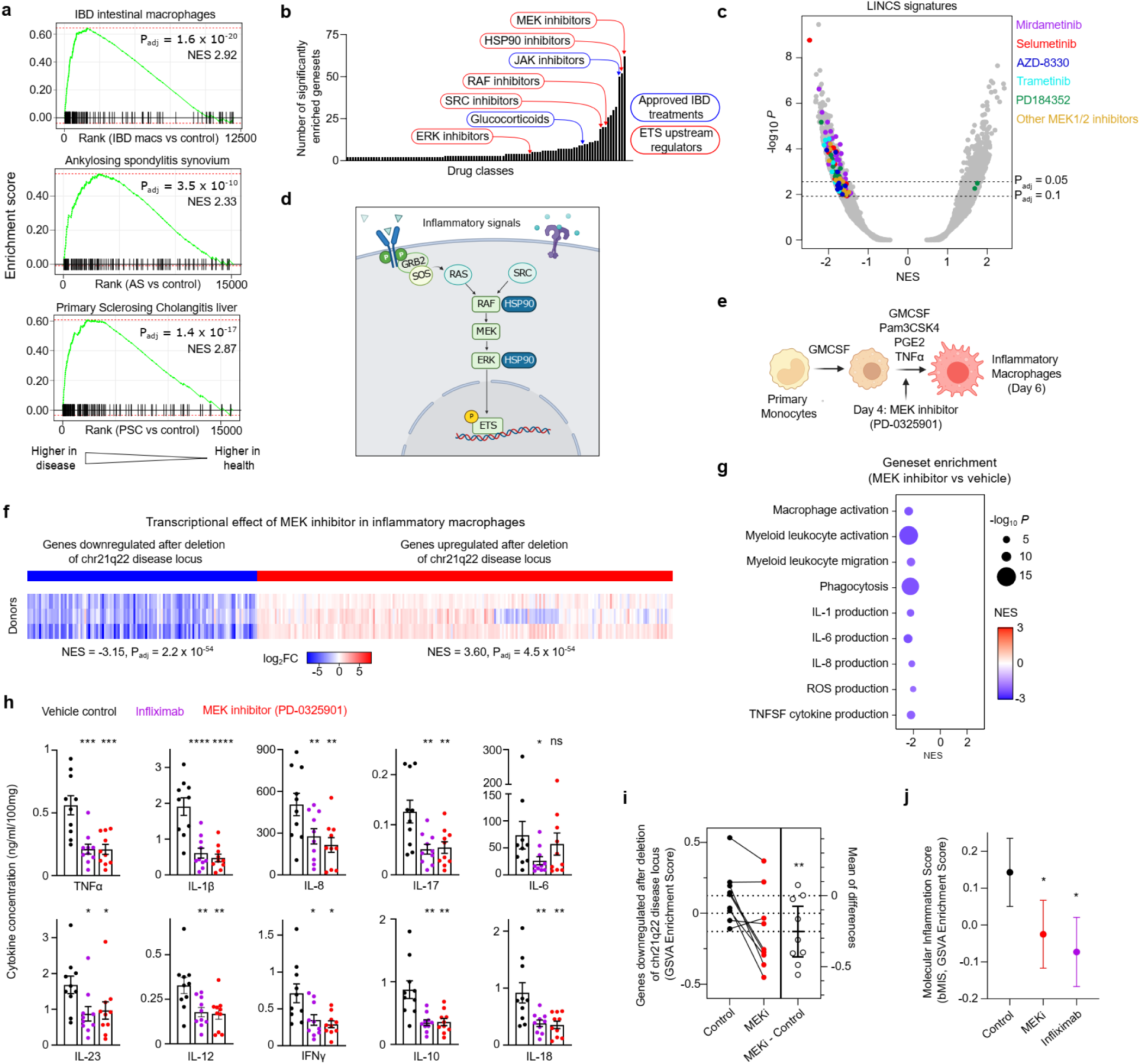
ETS2-driven inflammation is evident in disease and can be therapeutically targeted. **a.** Enrichment of chr21q22-regulated genes in IBD intestinal macrophages (top), AS synovium (middle), and PSC liver (bottom). All relative to unaffected tissue. Gene set comprised significantly downregulated genes following chr21q22 deletion. **b.** Candidate drug classes (from NIH LINCS database) that phenocopy the transcriptional consequences of *ETS2* disruption. **c.** fGSEA results for NIH LINCS drug signatures (FDR *P* estimated using adaptive multi-level split Monte-Carlo scheme; NES, normalised enrichment score). Significantly enriched MEK inhibitor gene sets coloured by molecule. **d.** Schematic of known upstream regulators of *ETS*-family transcription factors. **e.** Schematic of experiment for treating inflammatory macrophages with a MEK inhibitor (PD-0325901) **f.** Heatmap of relative expression (log2 fold-change) of chr21q22-regulated genes in inflammatory macrophages following MEK inhibition (compared to vehicle control, n=3). **g.** fGSEA of differentially-expressed genes between MEK inhibitor-treated and control inflammatory macrophages. Results shown for Gene Ontology Biological Pathways that were negatively enriched following *ETS2* editing. Dot size represents *P*-value and colour denotes NES. **h.** Cytokine secretion from IBD mucosal biopsies cultured for 18 hours with vehicle control, PD-0325901, or infliximab. **i.** Estimation plot of GSVA enrichment scores for chr21q22-downregulated genes in IBD intestinal biopsies following MEK inhibition (MEKi). Error bars indicate 95%CI. **j.** GSVA enrichment scores of a biopsy-derived molecular inflammation score (bMIS). Data in **h** and **i** represent mean±SEM. Wilcoxon matched-pairs test, two-tailed, n=10 (h), n=9 (j). * *P* < 0.05, ** *P* < 0.01, *** *P* < 0.001, **** *P* < 0.0001.

## Discussion

Arguably the greatest challenge in modern genetics is to translate the success of GWAS into a better understanding of human disease. Here, we show how this can provide insights into basic immunology as well as disease mechanisms, with investigation of a single pleiotropic risk locus leading to the discovery of a master regulator of human macrophage responses and a key pathogenic pathway that is potentially druggable. Monocyte-derived macrophages play an important role in the pathogenesis of many inflammatory diseases^14^, producing cytokines that are often targeted therapeutically. Blocking individual cytokines, however, is typically only effective in 30-40% of patients^44^ and there is a growing realisation that modulating several cytokines at once may be a better approach^45^. Modulating ETS2 signalling via MEK1/2 inhibition has a broad effect on multiple inflammatory cytokines, including TNFα, IL-23, and IL-12 – all targets of existing therapies – and IL-1β which has previously been linked to treatment-refractory IBD^46^ but is not directly modulated by other small molecules (e.g. JAK inhibitors). However, systemic use of MEK inhibitors may not be an ideal strategy for treating chronic disease due to the physiological roles of MEK in other tissues. Indeed, use of MEK inhibitors for other conditions is currently limited by severe opthalmic, cardiac, and pulmonary toxicities^47^. Targeting ETS2 directly – for example using PROTACs or molecular glues – or selectively delivering a MEK inhibitor to inflammatory macrophages via antibody-drug conjugates could potentially overcome this toxicity, and provide a safer means of inhibiting ETS2-driven inflammation.

These findings emphasise the importance of studying disease mechanisms beyond simple changes in gene expression, especially since the overlap between the chr21q22 risk haplotype and a macrophage eQTL had been noted previously^5^ without any indication as to the importance of the downstream biology. Indeed, it is even possible that *ETS2* is involved in other human pathology, aside from diseases associated with the chr21q22 locus. For example, Down’s syndrome – caused by trisomy of chromosome 21 – was recently proposed to be a cytokinopathy^48^, with increased basal expression of multiple inflammatory cytokines, including several that are specifically upregulated following *ETS2* overexpression (e.g. IL-1β, TNFα, and IL-6). Whether the additional copy of the *ETS2* gene contributes to this phenotype is unknown, but warrants further study. Relatedly, it is interesting to consider why a polymorphism that increases susceptibility to multiple inflammatory diseases is so common (risk allele frequency ∼75% in Europeans and >90% in Africans, **Extended Data Fig.9**). One possibility is that enhanced macrophage effector responses might provide a selective advantage during infection, which would explain why the ancestral risk allele has not been negatively selected. Recent studies have not found evidence for a strong selective sweep at this locus within the past few thousand years^49, 50^, but more ancient selection – or balancing selection maintaining variation – cannot be excluded, especially since rs2836882 appears to be an exceptionally old SNP (conservatively estimated at >500,000 years old; **Extended Data Fig.9**) and was even polymorphic between Neanderthals and Denisovans (**Extended Data Fig.9**).

In summary, using an intergenic GWAS hit as a starting point, we have identified a druggable pathway that is both necessary and sufficient for human macrophage responses during chronic inflammation. Furthermore, we show how genetic dysregulation of this pathway – via perturbation of pioneer factor binding at a critical long-range enhancer – confers susceptibility to multiple inflammatory diseases. This highlights the considerable, yet largely untapped, opportunity to better understand disease biology that is presented by non-coding genetic associations.

## Methods

### Analysis of existing data relating to chr21q22

We used IBD GWAS summary statistics^3^ to perform multiple causal variant fine-mapping using susieR^51^ with reference minor allele and LD information calculated from 503 European samples from 1000 Genomes phase 3. All R analyses used v.4.2.1. Palindromic SNPs (A/T or C/G) and any SNPs that didn’t match by position or alleles were pruned before imputation using the *ssimp* equations reimplemented in R. This did not affect any candidate SNP at chr21q22. We obtained SuSiE fine mapping results for *ETS2* (with identifier ENSG00000157557 or ILMN_1720158) in monocyte datasets from the eQTL Catalog.

Colocalisation analysis was performed using coloc v5.2.0^52^ using a posterior probability of H4 (PP.H4.abf) > 0.5 to call colocalisation.

Raw H3K27ac ChIP-seq data from primary human immune cells were downloaded from Gene Expression Omnibus (GEO series GSE18927 and GSE96014) and processed as described previously^53^.

Processed promoter-capture Hi-C data^15^ from 17 primary immune cell-types were downloaded from OSF (https://osf.io/u8tzp).

### Monocyte purification and macrophage differentiation

Leukocyte cones from healthy donors were obtained from NHS Blood and Transplant (Cambridge Blood Donor Centre, Colindale Blood Centre, or Tooting Blood Donor Centre). Peripheral blood mononuclear cells (PBMCs) were isolated by density centrifugation (Histopaque 1077, Sigma) and monocytes were positively selected using CD14 Microbeads (Miltenyi Biotec). Macrophage differentiation was performed using conditions that model chronic inflammation (TPP)^17^: 3 days GM-CSF (50ng/ml, Peprotech) followed by 3 days GM-CSF, TNFα (50ng/ml, Peprotech), PGE_2_ (1μg/ml, Sigma Aldrich), and Pam_3_CSK4 (1μg/ml, Invivogen). All cultures were performed at 37°C, 5% CO_2_ in antibiotic-free RPMI1640 media containing 10% FBS, GlutaMAX, and MEM Non-Essential Amino Acids (all ThermoFisher). Cells were detached using Accutase (Biolegend).

### Identifying a model of chronic inflammatory macrophages

Human monocyte-derived macrophage gene expression data files (n=299) relating to 64 different polarisation conditions were downloaded from Gene Expression Omnibus (GSE47189) and quantile normalised. Data from biological replicates were summarised to the median value for every gene. Gene set variation analysis^54^ (using the *GSVA* package in R) was performed to identify the polarisation condition that most closely resembled CD14+ monocytes/macrophages from active IBD – using disease-associated lists of differentially-expressed^55^.

### CRISPR-Cas9 editing of primary human monocytes

gRNA sequences were designed using CRISPick (formerly GPP sgRNA Designer) and synthesised by IDT. gRNA sequences: chr21q22 5’ gRNA, CCUGGCUGCCUCGCGUUUCC; chr21q22 3’ gRNA, CCUCGUCCAACAGAGAGCAA; ETS2 gRNA1, CAGACACAGAAUUACCCCAA; ETS2 gRNA2, UUGCUGCACGGGGUUAACAA. Alt-R CRISPR-Cas9 Negative Control crRNA #1 (IDT) was used as a non-targeting control. Cas9-gRNA ribonucleoproteins were assembled as described previously^53^ and nucleofected into 5×10^6^ monocytes in 100μL nucleofection buffer (Human Monocyte Nucleofection Kit, Lonza) using a Nucleofector 2b (Lonza, program Y-001). After nucleofection, monocytes were immediately transferred into 5ml of pre-warmed culture media in a 6-well flat-bottomed plate, and differentiated into macrophages under TPP conditions. Editing efficiency was quantified by PCR amplification of the target region in extracted DNA (chr21q22_Fw primer, GGTGGGGAGAGTTCCAAAGG; chr21q22_Rv, TCACCCTTCACCTCTTTGCT; ETS2_g1_Fw, TCCTGAAGGTCCCATGAAAG; ETS2_g1_Rv, TCATTATGGCTCTGGGGTTC; ETS2_g2_Fw, GCGGCACATTCATATCACAC; ETS2_g2_Rv, GCAGAATACCCCAAGCAAAA). Editing efficiency at the chr21q22 locus was measured via quantification of amplified fragments (2100 Bioanalyzer, Agilent) as previously described^53^.

Editing efficiency for individual gRNAs was assessed using the Inference of CRISPR Edits tool^56^ (ICE, Synthego).

### PrimeFlow RNA Assay

RNA abundance was quantified by PrimeFlow (ThermoFisher) in chr21q22-edited and unedited (NTC) cells on days 0, 3, 4, 5, and 6 of TPP differentiation. Target probes specific for *ETS2* (Alexa Fluor 647), *BRWD1* (Alexa Fluor 568) and *PSMG1* (Alexa Fluor 568) were used according to the manufacturer’s instructions. Data were analysed using FlowJo v10 (BD Biosciences).

### MPRA

Overlapping oligonucleotides containing 114-nt of genomic sequence were designed to tile the region containing chr21q22 candidate SNPs (99% credible set) at 50bp intervals. Six technical replicates were designed for every genomic sequence, each tagged by a unique 11-nt barcode. Additional oligonucleotides were included to test the expression-modulating effect of every candidate SNP in the 99% credible set. Allelic constructs were designed as described previously^53^ and tagged by 30 unique 11-nt barcodes. Positive and negative controls were included as described previously^53^. 170-nt oligonucleotides were synthesised as part of a larger MPRA pool (Twist Biosciences) containing the 16-nt universal primer site ACTGGCCGCTTCACTG, 114-nt variable genomic sequence, KpnI and XbaI restriction sites (TGGACCTCTAGA), an 11-nt barcode, and the 17-nt universal primer site AGATCGGAAGAGCGTCG. Cloning into the MPRA vector was performed as described previously^53^. A suitable promoter for the MPRA vector (RSV) was identified by testing promoter activities in TPP macrophages. The MPRA vector library was nucleofected into TPP macrophages (5µg vector into 5×10^6^ cells) in 100μl nucleofection buffer (Human Macrophage Nucleofection Kit, Lonza) using a Nucleofector 2b (program Y-011). To ensure adequate barcode representation, a minimum of 2×10^7^ cells were nucleofected for every donor (n=8). After 24 hours, RNA was extracted and sequencing libraries were made from mRNA or DNA input vector as described previously^53^. Library pools (of 6 samples each) were sequenced on an Illumina HiSeq2500 high output flow-cell (50bp, single-end reads) and data were pre-processed as previously described^53^. To identify regions of enhancer activity, a paired t-test was performed to identify genomic sequences that enhanced transcription. A sliding window analysis (300-bp window) was then performed across all tiling sequences using the *les* package in R. Expression-modulating variants were identified using QuASAR-MPRA^57^, as described previously^53^.

### BaalChIP

Publicly-available PU.1 ChIP-seq datasets from human macrophages were downloaded from GEO, and BAM files were examined (using the IGV genome browser) to identify rs2836882 heterozygotes (i.e. files containing both A and G allele reads at chr21:40,466,570; hg19). Two suitable samples were identified (GSM1681423 and GSM1681429) which were used for a Bayesian analysis of allelic imbalances in PU.1 binding, with correction for biases introduced by overdispersion and biases towards the reference allele – implemented in the *BaalChIP* package^22^ in R.

### Allele-specific PU.1 ChIP-genotyping

A 100ml blood sample was taken from five healthy individuals who were heterozygous at rs2836882 (assessed via Taqman genotyping, ThermoFisher). All participants provided written informed consent. Ethical approval was provided by the London - Brent Regional Ethics Committee (REC: 21/LO/0682). Monocytes were isolated from PBMC using CD14 Microbeads (Miltenyi Biotec) and differentiated into inflammatory macrophages using TPP conditions^17^. Following differentiation, macrophages were detached using Accutase and cross-linked for 10 min in fresh media containing 1% formaldehyde. Cross-linking was quenched with glycine for 5 min (final concentration 0.125M). Nuclei preparation and shearing were performed as described previously^53^ with 10 cycles sonication (30s ON/30s OFF, Bioruptor Pico, Diagenode). PU.1 was immunoprecipitated overnight at 4°C using a polyclonal anti-PU.1 antibody (1:25; Cell Signaling) with the SimpleChIP Plus kit (Cell Signaling). The ratio of rs2836882 alleles in the PU.1-bound DNA was quantified in duplicate by TaqMan genotyping (assay C 2601507_20). A standard curve was generated using fixed ratios of geneblocks containing either the risk or non-risk allele (200-nt genomic sequence centred on rs2836882; Genewiz).

### PU.1 MPRA-ChIP-seq

The MPRA vector library was transfected into TPP macrophages from six healthy donors. Assessment of PU.1 binding to SNP alleles was performed as described previously^53^, with minimal sonication (to remove contaminants while minimising chromatin shearing). Immunoprecipitation was performed overnight at 4°C using a polyclonal anti-PU.1 antibody (1:25; Cell Signaling) with the SimpleChIP Plus kit (Cell Signaling). Sequencing libraries were prepared from isolated plasmids as for MPRA and sequencing on a MiSeq (50bp, single-end reads).

### H3K27ac ChIP-seq

TPP macrophages from two rs2836882 major allele homozygotes and two minor allele homozygotes were harvested, cross-linked, and quenched as described above. Donors were identified through the NIHR BioResource. H3K27ac ChIP-seq was performed as described previously^53^ using an anti-H3K27ac antibody (1:250, Abcam) or an isotype control (1:500, rabbit IgG, Abcam). Libraries were sequenced on a HiSeq4000 (50bp, single-end reads). Raw data were processed, QC’d, and analysed as described previously^53^ (see Code Availability).

### Assays of macrophage effector functions

#### Flow cytometry

Expression of myeloid markers was assessed by flow cytometry (BD LSRFortessa^TM^ X-20). Panel: CD11b PE/Dazzle 594 (BioLegend), CD14 evolve605 (ThermoFisher), CD16 PerCP (BioLegend), CD68 FITC (BioLegend), Live/Dead Fixable Aqua Dead Cell Stain (ThermoFisher), and Fc Receptor Blocking Reagent (Miltenyi). Data were analysed using FlowJo v10 (BD Biosciences).

#### Cytokine quantification

Supernatants were collected on day 6 of TPP macrophage culture and frozen. Cytokine concentrations were quantified in duplicate via electrochemiluminescence using U-PLEX assays (Meso Scale Diagnostics).

#### Phagocytosis

Phagocytosis was assessed using fluorescently-labelled Zymosan particles (Green Zymosan, Abcam) according to the manufacturer’s instructions. Cells were seeded at 10^5^ cells/well in 96-well round bottom plates. Cytochalasin D (10μg/ml, ThermoFisher), an inhibitor of cytoskeletal rearrangement, was used as a negative control. Phagocytosis was quantified via flow cytometry, and a phagocytosis index was calculated (proportion of positive cells multiplied by their mean fluorescence intensity).

#### Extracellular ROS production

Extracellular ROS production was quantified using a Diogenes Enhanced Superoxide Detection Kit (National Diagnostics) according to the manufacturer’s protocol. Cells were seeded at a density of 10^5^ cells/well and pre-stimulated with PMA (200ng/ml, Sigma Aldrich).

#### Western blotting

Western blotting was performed as described previously^58^ using the following primary antibodies: rabbit anti-gp91*phox*, rabbit anti-p22*phox* (both Santa Cruz), rabbit anti-C17ORF62/EROS (Atlas), rabbit anti-actin (Abcam). Secondary antibody was anti-rabbit IgG-horseradish peroxidase (Cell Signaling). Chemiluminescence was recorded on a ChemiDoc Touch imager (Bio-Rad) following incubation of the membrane with ECL (ThermoFisher) or SuperSignal West Pico PLUS (ThermoFisher) reagent.

### RNA sequencing

RNA was isolated from macrophage lysates (AllPrep DNA/RNA Micro Kit, Qiagen) and sequencing libraries prepared from 10ng RNA using the SMARTer Stranded Total RNA-Seq Kit v2 - Pico Input Mammalian (Takara) following the manufacturer’s instructions. Libraries were sequenced on a NextSeq2000 (50bp, PE reads: CRISPR-based loss-of-function, roxadustat and PD-0325901 experiments) or a NovaSeq6000 (100bp, PE reads: overexpression experiments). Reads were trimmed using Trim Galore (Phred score 24), filtered to remove reads < 20bp, and ribosomal reads were removed using the BBSplit function of BBMap (BBMap, sourceforge.net/projects/bbmap/) with the Human ribosomal DNA complete repeating unit (GenBank: U13369.1). Reads were aligned to the human genome (hg38) using HISAT2 (ref.^59^) and converted to BAM files, sorted and indexed using SAMtools^60^. Gene read counts were obtained with the featureCounts program^61^ from Rsubread using the GTF annotation file for human genome build GRCh38 (version 102). Differential expression analysis was performed in R using the *limma* package^62^ with the voom transformation and including donor as a covariate.

### Gene set enrichment analysis

GSEA was performed using the *fGSEA*^63^ package in R. Gene sets were either obtained from Gene Ontology Biological Pathways (downloaded from MSigDB), experimentally-derived based on differential expression analysis, or sourced from published literature^7^. Pathways shown in Figures 2-5 are: GO:0002274, GO:0042116, GO:0097529, GO:0006909, GO:0071706, GO:0032732, GO:0032755, GO:0032757, GO:2000379, GO:0009060, GO:0006119, and GO:0045649. Statistical significance was calculated using the adaptive multilevel split Monte Carlo method.

### *In vitro* transcription

The cDNA sequence for *ETS2* (NM005329.5) preceded by a Kozak sequence was synthesised and cloned into a TOPO vector. This was linearised and a PCR amplicon of the *ETS2* gene generated, adding a T7 promoter and an AG initiation sequence (Phusion, NEB): Fw primer: GCTAATACGACTCACTATAAGGACAGGCCACCATGAATGATTTCGGAATC, Rv primer: TCAGTCCTCCGTGTCGG). A reverse complement (control) amplicon was also generated: Fw primer: GCTAATACGACTCACTATAAGGACAGGCCACCTCAGTCCTCCGTGTCGG, Rv primer: GCCACCATGAATGATTTCGGAATC). These amplicons were used as templates for *in vitro* transcription using the HiScribe T7 mRNA Kit with CleanCap® Reagent AG kit (NEB) according to the manufacturer’s instructions, but with substitution of N1-methyl-pseudouridine for uridine and methylcytidine for cytidine (both Stratech) to minimise non-specific cellular activation by the transfected mRNA. mRNA was purified using a MEGAclear Transcription Clean-Up Kit (ThermoFisher) and polyadenylated using an E. coli Poly(A) Polymerase (NEB) before further clean-up (MEGAclear), quantification and analysis of product size (NorthernMax®-Gly gel, ThermoFisher). For optimising overexpression conditions, GFP mRNA was produced using the same method: Fw primer GCTAATACGACTCACTATAAGGACAGGCCACCATGGTGAGCAAGGGCGAG, Rv primer TTACTTGTACAGCTCGTCCATGC).

### mRNA overexpression

Lipofectamine MessengerMAX (ThermoFisher) was diluted in Opti-MEM (1:75 v/v), vortexed and incubated at room temperature for 10 minutes. IVT mRNA was then diluted in a fixed volume of Opti-MEM (112.5µl per transfection), mixed with an equal volume of diluted Lipofectamine MessengerMAX and incubated for a further 5 minutes at room temperature. The transfection mix was then added dropwise to 2.5×10^6^ M0 macrophages (pre-cultured for 6 days in a 6-well plate in antibiotic-free RPMI1640 macrophage media containing M-CSF (50ng/ml, Peprotech) – with media change on d3). For GFP overexpression, cells were detached using Accutase 18 hours after transfection and GFP expression was measured by flow cytometry. For ETS2 / control overexpression, either 250ng or 500ng mRNA was transfected and low dose LPS (0.5ng/ml) was added 18 hours after transfection, and cells detached using Accutase 6 hours later (n=8 donors). Representative *ETS2* expression in untransfected macrophages obtained from previous data (GSE193336).

### SNPsea

Pathway analysis of 241 IBD-associated GWAS hits^3^ was performed using SNPsea^34^. In brief, linkage intervals were defined for every lead SNP based on the furthest correlated SNPs (r^2^ > 0.5 in 1000 Genomes, EUR population) and extended to the nearest recombination hotspots with recombination rate >3 cM/Mb. If no genes were present in this region, the linkage interval was extended up- and down-stream by 500kb. Genes within linkage intervals were tested for enrichment within 7,660 pathways, comprising 7,658 Gene Ontology Biological Pathways and 2 lists of ETS2-regulated genes (either those significantly downregulated following *ETS2* disruption with gRNA1 or those significantly upregulated following ETS2 overexpression, based on a consensus list obtained from differential expression analysis including all samples and using donor and mRNA quantity as covariates). The analysis was performed using a single score mode: assuming that only one gene per linkage interval is associated with the pathway. A null distribution of scores for each pathway was performed by sampling random SNP sets matched on the number of linked genes (5,000,000 iterations). A permutation *P-* value was calculated by comparing the enrichment of the IBD-associated gene list with the null distribution. Gene sets relating to the following IBD-associated pathways were extracted for comparison: NOD2 signalling (GO:0032495), Integrin signalling (GO:0033627, GO:0033622), TNFα signalling (GO:0033209, GO:0034612), Intestinal epithelium (GO:0060729, GO:0030277), Th17 cells (GO:0072539, GO:0072538, GO:2000318), T cell activation (GO:0046631, GO:0002827), IL-10 signalling (GO:0032613, GO:0032733), and autophagy (GO:0061919, GO:0010506, GO:0010508, GO:1905037, GO:0010507).

### ETS2 co-expression

Genes co-expressed with ETS2 across 64 human monocyte-derived macrophage polarisation conditions (normalised data from GSE47189) were identified using the rcorr function in the *Hmisc* package in R.

### 13C-glucose GC-MS

*ETS2*-edited or unedited TPP macrophages were generated in triplicate for each donor and on day 6, media was removed, cells washed with PBS, and new media with labelled glucose was added. Labelled media: RPMI1640 Medium, no glucose (ThermoFisher); 10% FBS (ThermoFisher); GlutaMAX (ThermoFisher); ^13^C-labelled glucose (Cambridge Isotype Laboratories). After 24 hours – a time point selected from a time-course to establish steady-state conditions – supernatants were snap-frozen and macrophages detached by scraping. Macrophages were washed three times with ice-cold PBS, counted, re-suspended in 600µl ice-cold chloroform:methanol (2:1, v/v) and sonicated in a waterbath (3 x 8 mins). All extraction steps were performed at 4°C as previously described^64^. Samples were analysed in an Agilent 7890B-7000C GC–MS system. Spitless injection (injection temperature 270°C) onto a DB-5MS (Agilent) was used, using helium as the carrier gas, in electron ionization mode. The initial oven temperature was 70°C (2 min), followed by temperature gradients to 295 °C at 12.5 °C per min and to 320 °C at 25 °C per min (held for 3 min). Scan range was m/z 50-550. Data analysis was performed using in-house software MANIC (version 3.0), based on the software package GAVIN^65^. Label incorporation was calculated by subtracting the natural abundance of stable isotopes from the observed amounts. Total metabolite abundance was normalised to the internal standard (scyllo-inositol^64^).

### Roxadustat

*ETS2-*edited or unedited TPP macrophages were generated as described previously. On day 5 of culture, cells were detached (Accutase) and re-plated at a density of 10^5^ cells/well in 96-well round bottom plates in TPP media containing Roxadustat (FG-4592, 30μM). After 12 hours, cells were harvested for functional assays and RNA-seq as described.

## CUT&RUN

Pre-cultured TPP macrophages were harvested and processed immediately using the CUT&RUN Assay kit (Cell Signaling) according to the manufacturer’s instructions but omitting the use of ConA-coated beads. In brief, 5×10^5^ cells per reaction were pelleted, washed, and resuspended in Antibody Binding buffer. Cells were incubated with antibodies: anti-ETS2 (1:100, ThermoFisher) or IgG control (1:20, Cell Signaling) for 2h at 4°C. After washing in Digitonin Buffer, cells were incubated with pA/G-MNase for 1h at 4°C. Cells were washed twice in Digitonin Buffer, resuspended in the same buffer and cooled for 5 minutes on ice. Calcium chloride was added to activate pA/G-MNase digestion (30 min, 4°C) before the reaction was stopped and cells incubated at 37°C for 10 min to release cleaved chromatin fragments. Supernatants were collected by centrifugation and DNA extracted using spin columns (Cell Signaling). Library preparation was performed using a protocols.IO protocol (dx.doi.org/10.17504/protocols.io.bagaibse) with the NEBNext Ultra II DNA Library Prep Kit. Size selection was performed using AMPure XP beads (Beckman Coulter) and fragment sizes assessed using an Agilent 2100 Bioanalyzer (High Sensitivity DNA kit). Equimolar pools of indexed libraries were sequenced on a NovaSeq6000 (100bp PE reads). Raw data were analysed using guidelines from the Henikoff lab^66^. Briefly, paired-end reads were trimmed using Trim Galore and aligned to the human genome (GRCh37/hg19) using Bowtie2. BAM files were sorted, merged (technical and, where indicated, biological replicates), re-sorted and indexed using SAMtools. Picard was used to mark unmapped reads and SAMtools to remove these reads, re-sort and re-index. Bigwig files were created using the deepTools bamCoverage function. Processed data were initially analysed using the nf-core CUT&RUN pipeline v3.0, using CPM normalisation and default MACS2 parameters for peak calling. This analysis yielded acceptable quality metrics (including an average FRiP score of 0.23) but there were a high number of peaks with low fold enrichment (<4) over control. We therefore applied more stringent parameters for peak calling (--qvalue 0.05 -f BAMPE --keep-dup all -B --nomodel) and applied an irreproducible discovery rate (IDR; cut-off 0.001) to identify consistent peaks between replicates – implemented in the *idr* package in R (code: https://github.com/JamesLeeLab/chr21q22_manuscript/CUT&RUN/CUTRUN_pipeline.sh). Enrichment of an ETS2 binding motif in consensus IDR peaks was calculated using TFmotifView^67^ using global genomic controls. Overlap between consensus IDR peaks and the core promoter (-250bp to +35bp from TSS) and/or putative cis-regulatory elements of ETS2-regulated genes was assessed using lists of differentially-expressed genes following *ETS2* disruption with gRNA1 or *ETS2* overexpression (based on a consensus across mRNA doses, as described earlier). Putative cis-regulatory elements were defined as shared interactions (CHiCAGO score > 5) in monocyte, M0 and M1 macrophage samples from publicly-available promoter-capture Hi-C data^15^.

### ATAC-seq

ATAC-seq in TPP macrophages was performed using the Omni-ATAC protocol^68^ with the following modifications: cell number was increased to 75,000 cells, cell lysis time was increased to 5 minutes; volume of Tn5 transposase in the transposition mixture was doubled; duration of the transposition step was extended to 40 minutes. Amplified libraries were cleaned using AMPure XP beads (Beckman Coulter) and sequenced on a NovaSeq6000 (100bp PE reads). Data were processed as described previously^69^.

### Chr21q22 disease datasets

Publicly-available raw RNA-seq data from the affected tissues of chr21q22-associated diseases (and controls from the same experiment) were downloaded from GEO: IBD macrophages (GSE123141), primary sclerosing cholangitis liver (GSE159676), ankylosing spondylitis synovium (GSE41038). Reads were trimmed, filtered, and aligned as described earlier. For each disease dataset, a ranked list of genes was obtained by differential expression analysis between cases and controls using *limma* with voom transformation. For IBD macrophages, only IBD samples with active disease were included. fGSEA using ETS2-regulated gene lists was performed as described.

### LINCS signatures

31,027 lists of down-regulated genes following exposure of a cell line to a small molecule were obtained from the NIH LINCS database^8^ (downloaded in January 2021). These were used as gene sets for fGSEA (as described) with a ranked list of genes obtained by differential expression analysis between *ETS2*-edited and unedited TPP macrophages (gRNA1) – using *limma* with voom transformation and donor as a covariate. Drug classes for gene sets with FDR *P* < 0.05 were manually assigned based on known mechanisms of action.

### PD-0325901

TPP macrophages were generated as described previously. On day 4 of culture, PD-0325901 (0.5μM, Sigma) or vehicle (DMSO) were added. Cells were harvested on day 6 and RNA was extracted and sequenced as described.

### Colonic biopsies

During colonoscopy, intestinal mucosal biopsies (6 per donor) were collected from 10 IBD patients (7 ulcerative colitis, 3 Crohn’s disease). All had endoscopically active disease and were not receiving immunosuppressive or biologic therapies. All biopsies were collected from a single inflamed site. All patients provided written informed consent. Ethical approval was provided by the London - Brent Regional Ethics Committee (REC: 21/LO/0682). Biopsies were collected into Opti-MEM and within 1 hour were weighed and placed in pairs onto a transwell insert (ThermoFisher) – designed to create an air-liquid interface^70^ – in a 24-well plate. Each well contained 1ml media and was supplemented with either DMSO (vehicle control), PD-0325901 (0.5μM) or infliximab (10μg/ml; MSD). Media: Opti-MEM I (Gibco); GlutaMAX (ThermoFisher); 10% FBS (ThermoFisher); MEM Non-Essential Amino Acids (ThermoFisher); 1% sodium pyruvate (ThermoFisher); 1% penicillin/streptomycin (ThermoFisher); 50μg/ml gentamicin (Merck). After 18 hours, supernatants and biopsies were snap frozen. Supernatant cytokine concentrations were quantified using LEGENDplex Human Inflammation Panel (Biolegend). RNA was extracted from biopsies and libraries prepared as described earlier (n=9, RNA from one donor was too degraded). Sequencing was performed on a NovaSeq 6000 (100bp, PE reads). Data were processed as described earlier and GSVA was performed for ETS2-regulated genes and biopsy-derived signatures of IBD-associated inflammation^43^.

### Chr21q22 genotypes in archaic humans

Using publicly available genomes from seven Neanderthal individuals^71–74^, one Denisovan individual^75^, and one Neanderthal and Denisovan F1 individual^76^, we called genotypes at the disease-associated chr21q22 candidate SNPs from the respective BAM files using “bcftools mpileup” with base and mapping quality options -q 20 -Q 20 -C 50 and using “bcftools call -m -C alleles”, specifying the two alleles expected at each site in a targets file (-T option). From the resulting vcf file, we extracted the number of reads supporting the reference and alternative alleles stored in the “DP4” field.

### Inference of Relate genealogy at rs2836882

We used genome-wide genealogies previously inferred for samples of the Simons Genome Diversity Project^77^ dataset (https://reichdata.hms.harvard.edu/pub/datasets/sgdp/) using Relate^78, 79^. These genealogies were downloaded from https://www.dropbox.com/sh/2gjyxe3kqzh932o/AAAQcipCHnySgEB873t9EQjNa?dl=0. Using the inferred genealogies, the genealogy at rs2836882 (chr21:40466570) was plotted using the TreeView module of Relate.

### Statistical methodology

Statistical methods used in MPRA analysis, fGSEA, and SNPsea are described above. For other analyses, comparison of continuous variables between paired samples in two groups was performed using a Wilcoxon matched-pairs test for non-parametric data or a paired t-test for parametric data. Comparison against a hypothetical value was performed using a Wilcoxon signed-rank test for non-parametric data or one sample t-test for parametric data. A Shapiro-Wilk test was used to confirm normality. Two-tailed tests were used as standard unless a specific hypothesis was being tested. Sample sizes are provided in respective sections.

## Code availability

Code to reproduce analyses are available at https://github.com/JamesLeeLab/chr21q22_manuscript and https://github.com/chr1swallace/ibd-ets2-analysis.

## Data availability

The datasets produced in this study, including raw and processed files, have been uploaded to the following databases and will be made publicly available in advance of publication.

Gene Expression Omnibus: MPRA (GSE229472), RNA-seq of *ETS2* or chr21q22-edited TPP macrophages (GSE229569), RNA-seq of *ETS2* overexpression (GSE229744), RNA-seq of MEK inhibitor-treated TPP macrophages (GSE229743), H3K27ac ChIP-seq in TPP macrophages (GSE229464), ATAC-seq in TPP macrophages (GSE229624), ETS2 CUT&RUN (GSE229745), biopsy RNA-seq data (GSE230020).

MetaboLights: Metabolomics (MTBLS7665).

## Acknowledgements

We thank members of the Lee lab and Arthur Kaser for helpful discussions and Gitta Stockinger, Carola Vinuesa, Charlie Swanton, Rickie Patani, and Caetano Reis e Sousa for critical reading of the manuscript. We thank Chris Cheshire, the Francis Crick Institute Advanced Sequencing Facility and Flow Cytometry STP for technical support, Laura Lucaciu for help with patient recruitment, and Sarah Edwards for providing infliximab. We thank NIHR BioResource volunteers for their participation, and acknowledge NIHR BioResource centres, NHS Blood and Transplant, and NHS staff for their contribution. This work was supported by Crohn’s and Colitis UK (M2018-3), the Wellcome Trust (Sir Henry Wellcome Fellowship to L.S.: 220457/Z/20/Z, Investigator Award to P.S.: 217223/Z/19/Z, Senior Fellowship to C.W.: WT220788, Clinical Research Career Development Fellowship to M.Z.C.: 222056/Z/20/Z, Wellcome-Beit Prize Clinical Career Development Fellowship to D.C.T.: 206617/A/17/A, and Intermediate Clinical Fellowship to J.C.L.: 105920/Z/14/Z), and the Francis Crick Institute, which receives its core funding from Cancer Research UK (CC2219, FC001595), the UK Medical Research Council (CC2219, FC001595), and the Wellcome Trust (CC2219, FC001595). P.S. is additionally supported by the European Molecular Biology Organisation, the Vallee Foundation, and the European Research Council (852558). C.W. is additionally supported by the Medical Research Council (MC UU 00002/4), GSK, MSD, and the NIHR Cambridge BRC (BRC-1215-20014). D.C.T. is additionally supported by the Sidharth Burman endowment, and J.C.L. is a Lister Institute Prize Fellow. The funders had no role in study design, data collection and analysis, decision to publish, or preparation of the manuscript. Experimental schematics in Figs. 1b, 2a, 3a, 5e created using BioRender. For the purpose of Open Access, the authors have applied a CC BY public copyright licence to any Author Accepted Manuscript version arising from this submission.

## Author contributions

Conceptualisation, J.I.M., P.S., M.Z.C., C.W., D.C.T., J.C.L.; Methodology, C.T.S., C.B., M.S.D.S., L.O.R., L.S., J.I.M., C.W., J.C.L.; Software, C.B., M.S.D.S., L.S., J.I.M., C.W.; Investigation, C.T.S., C.B., T.T.S., A.P.P., C.P.J., I.P., M.S.D.S., L.O.R., L.S., E.C.P., W.E., A.P.R., C.D.M., C.W, J.C.L.; Resources, C.T.S., C.B., M.S.D.S., J.C.L.; Formal analysis, C.T.S., C.B., M.S.D.S., L.S., C.W., J.C.L.; Writing – Original Draft, C.T.S., C.B., J.C.L.; Writing – Review & Editing, all authors; Funding Acquisition, J.C.L.; Supervision, J.I.M., P.S., C.W., D.C.T., J.C.L.

## Additional information

**Supplementary information:** results of differential expression analysis in *ETS2*-edited or *ETS2* overexpression experiments are available in Supplementary Tables 1 and 2.

**Correspondence and requests for materials** should be addressed to James C. Lee.

