## Extended Data for "A disease-associated gene desert orchestrates macrophage inflammatory responses via ETS2"

**Extended Data – contents:**

- Extended Data Figure 1. Disease-associated variation at chr21q22 correlates with *ETS2* expression in monocytes.
- Extended Data Figure 2. CRISPR-Cas9 editing of the chr21q22 locus and *ETS2* in monocytes.
- Extended Data Figure 3. Optimisation of MPRA and mRNA overexpression in primary human macrophages.
- Extended Data Figure 4. Functional consequences of allelic variation at rs2836882.
- Extended Data Figure 5. Deletion of the chr21q22 disease-associated enhancer phenocopies *ETS2* disruption.
- Extended Data Figure 6. Metabolic effects of *ETS2* disruption.
- Extended Data Figure 7. The transcriptional signature of *ETS2* is detectable in affected tissues from chr21q22-linked diseases.
- Extended Data Figure 8. Effect of MEK1/2 inhibition on *ETS2*-regulated genes.
- Extended Data Figure 9. Geographic distribution and history of rs2836882.
- Extended Data Table 1. IBD risk genes downregulated following ETS2 disruption.

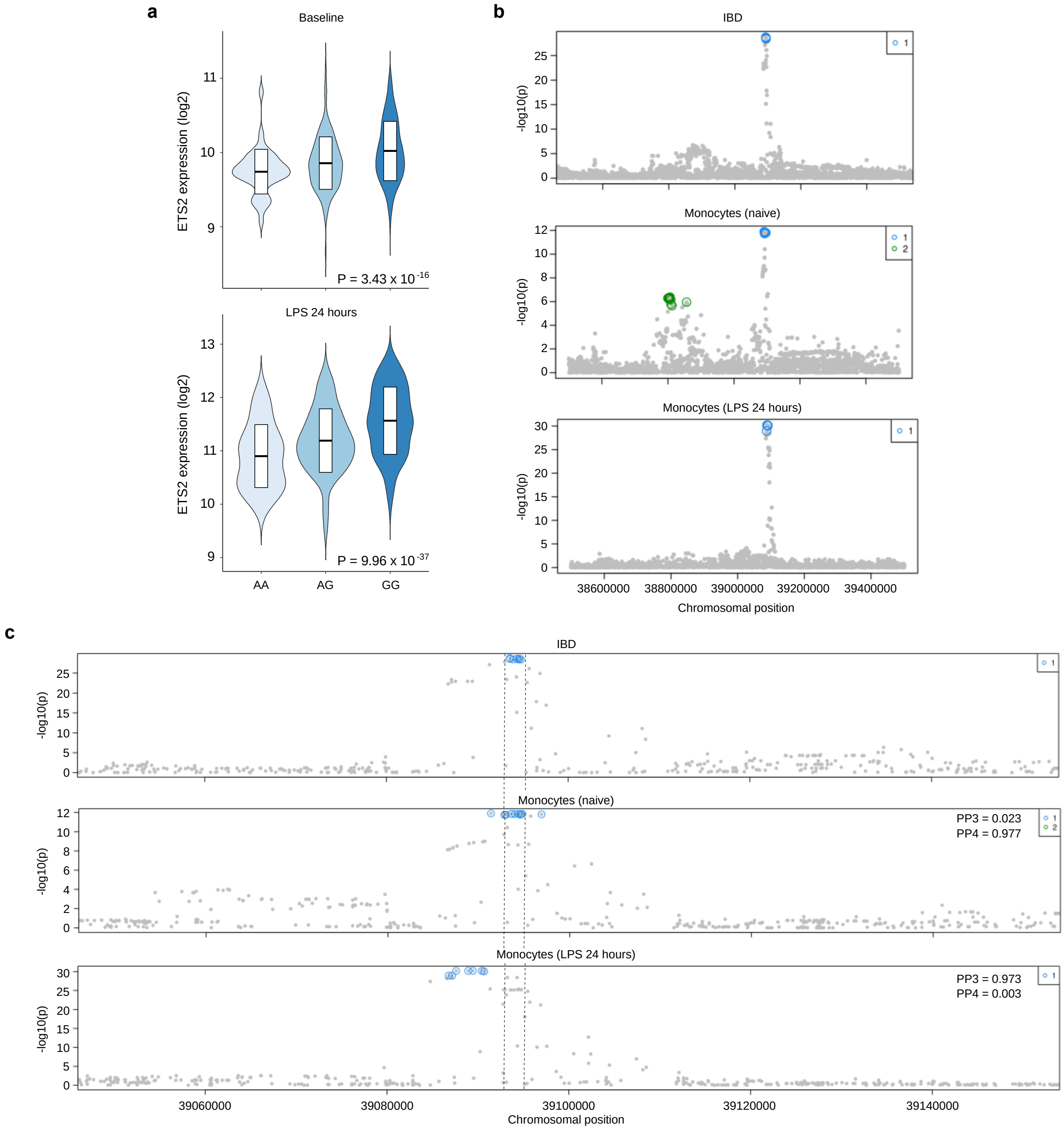


**Extended Data Figure 1. Disease-associated variation at chr21q22 correlates with *ETS2* expression in monocytes.**

**a.** Violin plots depicting *ETS2* expression stratified by rs2836882 genotype in resting (upper panel) and activated monocytes (lower panel). Inset boxes show median±IQR; data from ref.16. **b.** Manhattan plots depicting the associations at chr21q22 with IBD (upper panel), *ETS2* expression in resting monocytes (middle panel) and *ETS2* expression in LPS-stimulated monocytes (lower panel). Coloured circles denote independent signals at the locus. Genome coordinates hg38. **c.** Co-localisation analysis to determine the posterior probability that a shared causal variant underlies the chr21q22 IBD association and the *ETS2* eQTL in resting or activated monocytes. PP3, probability that GWAS and eQTL signals are independent. PP4, probability that a shared variant is responsible for both associations.

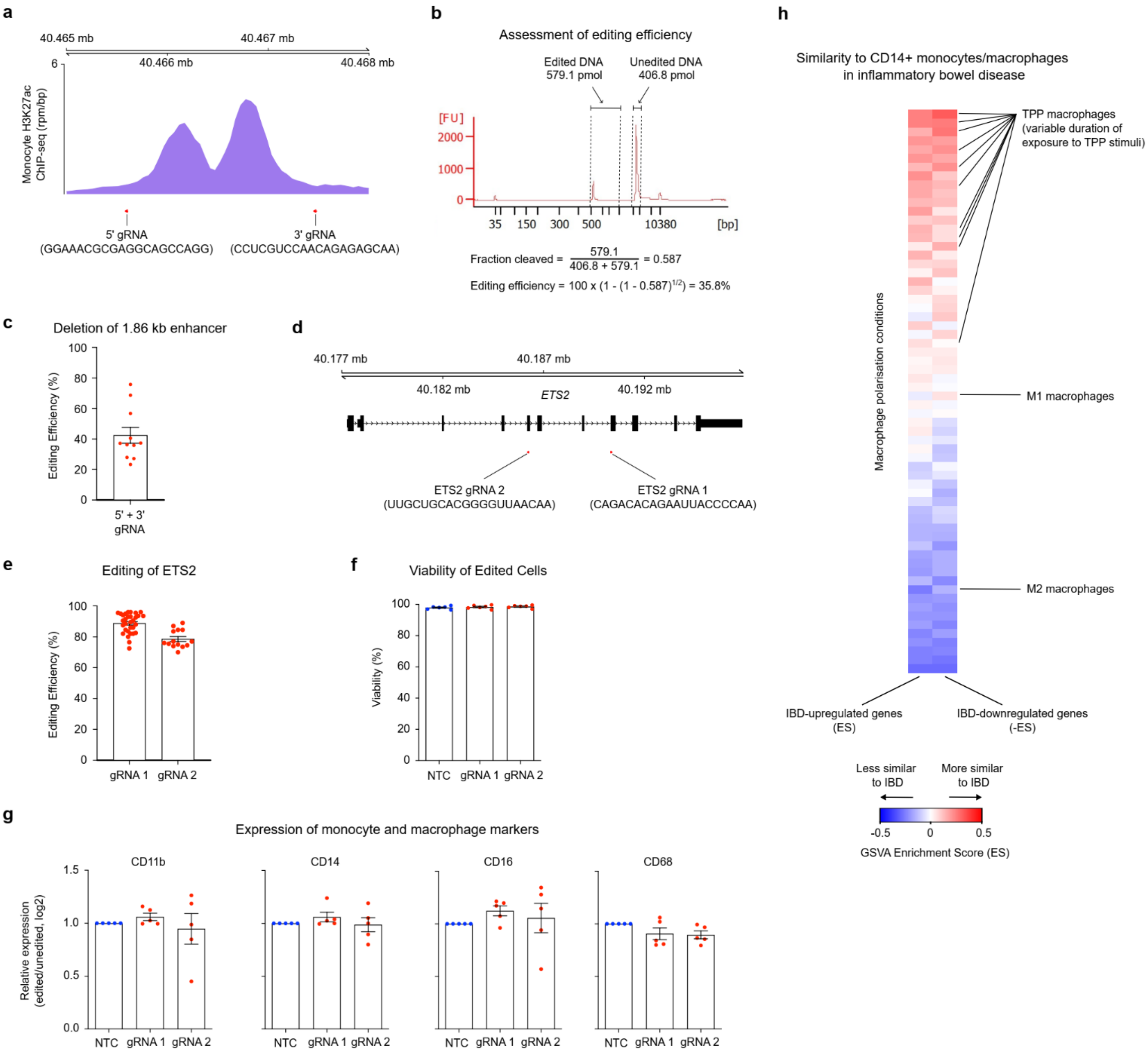


**Extended Data Figure 2. CRISPR-Cas9 editing of the chr21q22 locus and *ETS2* in monocytes.**

**a.** Cas9 gRNAs were designed to flank the enhancer region within the chr21q22 locus at the indicated sites. Lower plot shows H3K27ac ChIP-seq data in primary human monocytes. **b.** Representative bioanalyzer trace of PCR amplicon from target region following monocyte transfection with an equimolar mix of Cas9 RNPs containing 5’ or 3’ chr21q22 gRNAs. Method for calculating editing efficiency shown with example calculation. **c.** Editing efficiency at the chr21q22 locus. Mean frequency of enhancer deletion: 42.4%. **d.** Location and sequence of gRNAs used to disrupt *ETS2*. **e.** Editing efficiency at the *ETS2* gene. Mean editing efficiency: gRNA1, 89.7%; gRNA2, 78.6%. **f.** Viability following monocyte nucleofection with Cas9 RNPs and macrophage differentiation. Mean viabilities: NTC, 97.9%; gRNA1: 98.3%; gRNA2, 98.6%. **g.** Expression of myeloid lineage markers following *ETS2* editing and TPP differentiation. **h.** Gene set variance analysis in 64 different macrophage polarisation conditions to identify enrichment of gene sets significantly upregulated or downregulated in CD14+ monocytes/macrophages from IBD patients (compared to healthy controls). Error bars represent mean±SEM.

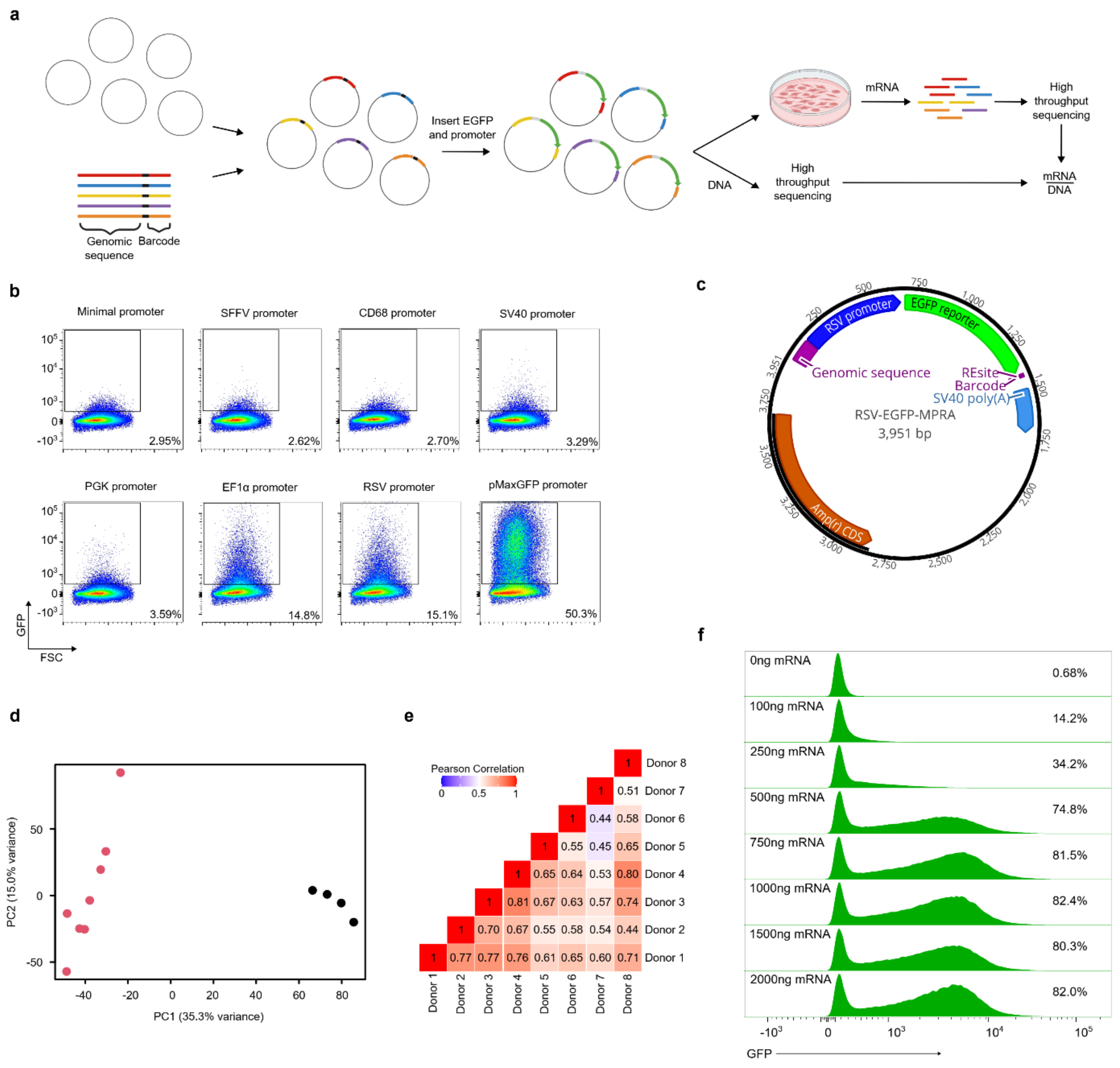


**Extended Data Figure 3. Optimisation of MPRA and mRNA overexpression in primary human macrophages.**

**a.** Schematic of MPRA. A library of oligonucleotides (each containing a genomic sequence and unique barcode, separated by restriction sites) is cloned into a pGL4.10M cloning vector. A promoter and reporter gene are then inserted using directional cloning. The resulting plasmids are transfected into primary human macrophages (TPP) and RNA is extracted after 24 hours. Barcode abundance in cellular mRNA and input DNA library are quantified by high-throughput sequencing, and mRNA barcode counts are normalised to corresponding counts in DNA library to assess expression-modulating activity. **b.** Identification of suitable promoters for MPRA in TPP macrophages. TPP macrophages were transfected with a reporter vectors, each with GFP expression under the control of a different promoter. GFP expression was quantified by flow cytometry after 24 hours. **c.** Adapted MPRA vector for use in primary human macrophages, containing RSV promoter. **d.** Principal component analysis of element counts (sum of barcodes tagging same genomic sequence) in mRNA from TPP macrophages (n=8 donors; red) and four replicates of DNA vector (black). **e.** Heatmap showing pairwise correlation of expression-modulating activity of all constructs between donors. **f.** Primary human macrophages (M0) were transfected with different quantities of GFP mRNA using Lipofectamine MessengerMAX. GFP expression was quantified by flow cytometry 18 hours after transfection.

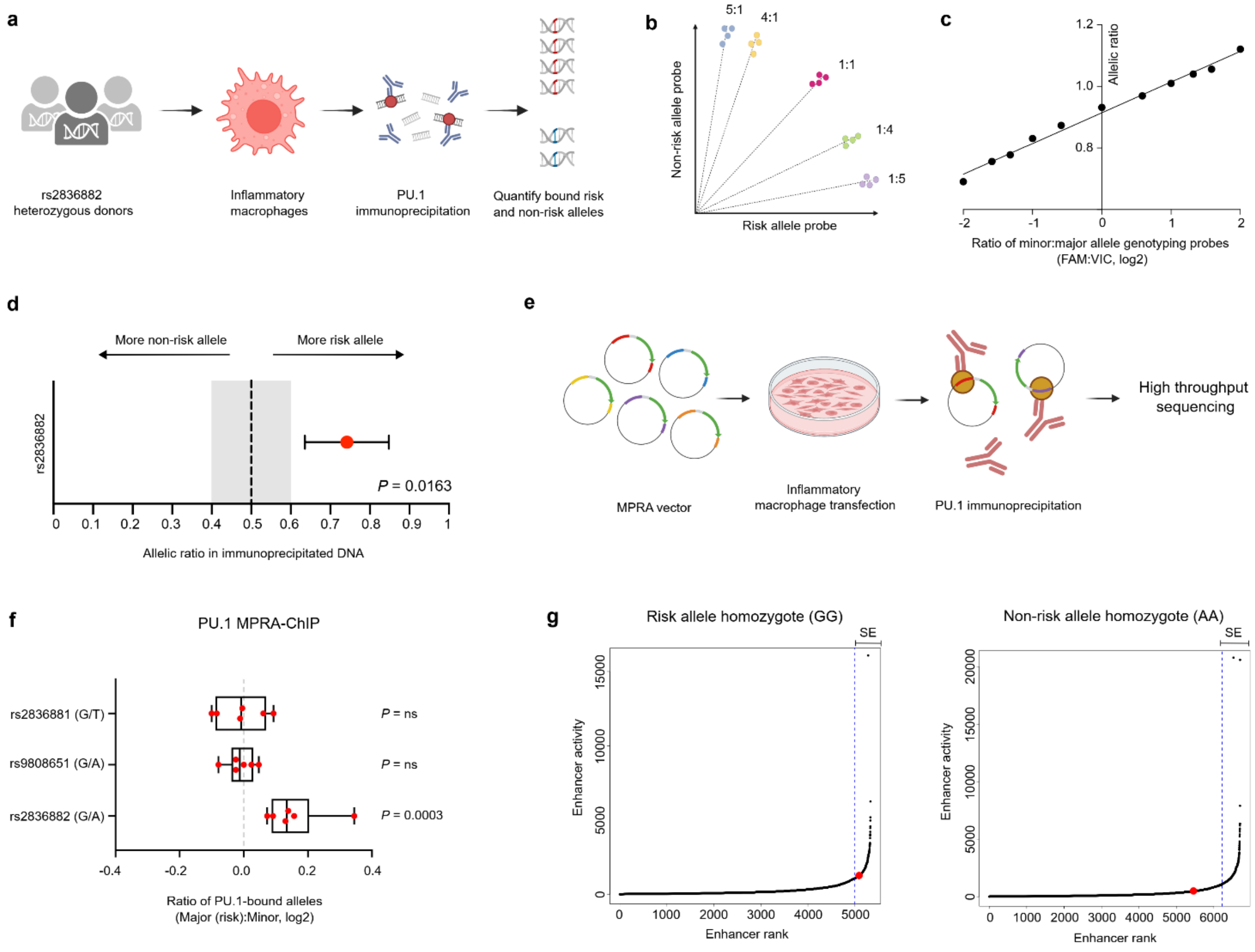


**Extended Data Figure 4. Functional consequences of allelic variation at rs2836882.**

**a.** Schematic of PU.1 ChIP-genotyping assay to assess allele-specific PU.1 binding at rs2836882 in human macrophages. **b.** Schematic of standard curve generation by TaqMan genotyping various pre-defined ratios of risk and non-risk containing DNA sequences. **c.** Standard curve generated using different allelic ratios of 200-nt DNA geneblocks centred on either the major (risk) or minor (non-risk) rs2836882 allele. **d.** Allele-specific PU.1 binding at rs2836882 in TPP macrophages (one-sample *t*-test, two-tailed, n=5). Error bars represent 95% confidence interval. **e**. Schematic of PU.1 MPRA-ChIP assay to assess allele-specific PU.1 binding at individual SNPs within chr21q22 enhancer. **f**. Allele-specific PU.1 binding at SNPs within chr21q22 enhancer in TPP macrophages. Data represents the allelic ratio of normalised PU.1 binding for constructs centred on the SNP allele from the MPRA library (fixed-effects meta-analysis of QuASAR-MPRA results, two-tailed, n=6). **g.** Rank Ordering of Super-Enhancers (ROSE) analysis of H3K27ac ChIP-seq data from TPP macrophages from major (left) and minor (right) allele homozygotes. Dashed line denotes inflection point of curve, with enhancers above this point being denoted as super-enhancers. Red points indicate rs2836882-containing chr21q22 enhancer. SE, super-enhancer.

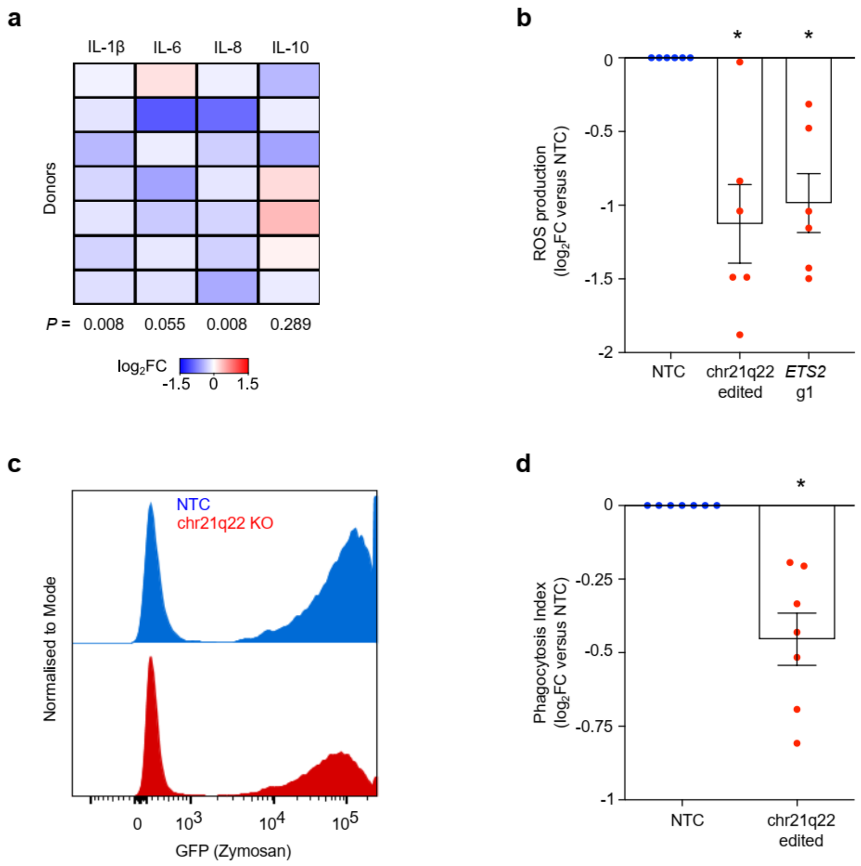


**Extended Data Figure 5. Deletion of the chr21q22 disease-associated enhancer phenocopies *ETS2* disruption.**

**a.** Cytokine secretion from TPP macrophages following deletion of the chr21q22 enhancer. Heatmap shows log2 fold-change of cytokine concentrations in the supernatants of chr21q22-edited inflammatory macrophages relative to non-targeting control (NTC) cells (Wilcoxon signed rank test, one-tailed). **b.** Extracellular ROS production by unedited (NTC), chr21q22-edited, and *ETS2* g1-edited TPP macrophages, quantified using a chemiluminescence assay. Points represent log2 fold-change of area under curve (AUC) for edited versus unedited cells (Wilcoxon signed-rank test). **c.** Representative flow cytometry histogram demonstrating phagocytosis of fluorescently-labelled zymosan particles by chr21q22-edited and unedited (NTC) TPP macrophages. **d.** Phagocytosis index for unedited (NTC) and chr21q22-edited TPP macrophages, calculated as proportion of positive cells multiplied by mean fluorescence intensity of positive cells (488 nm channel). Plot shows log2 fold-change of phagocytosis index for chr21q22-edited cells relative to unedited cells from same donor (Wilcoxon signed-rank test). Error bars represent mean±SEM. * $P < 0.05$.

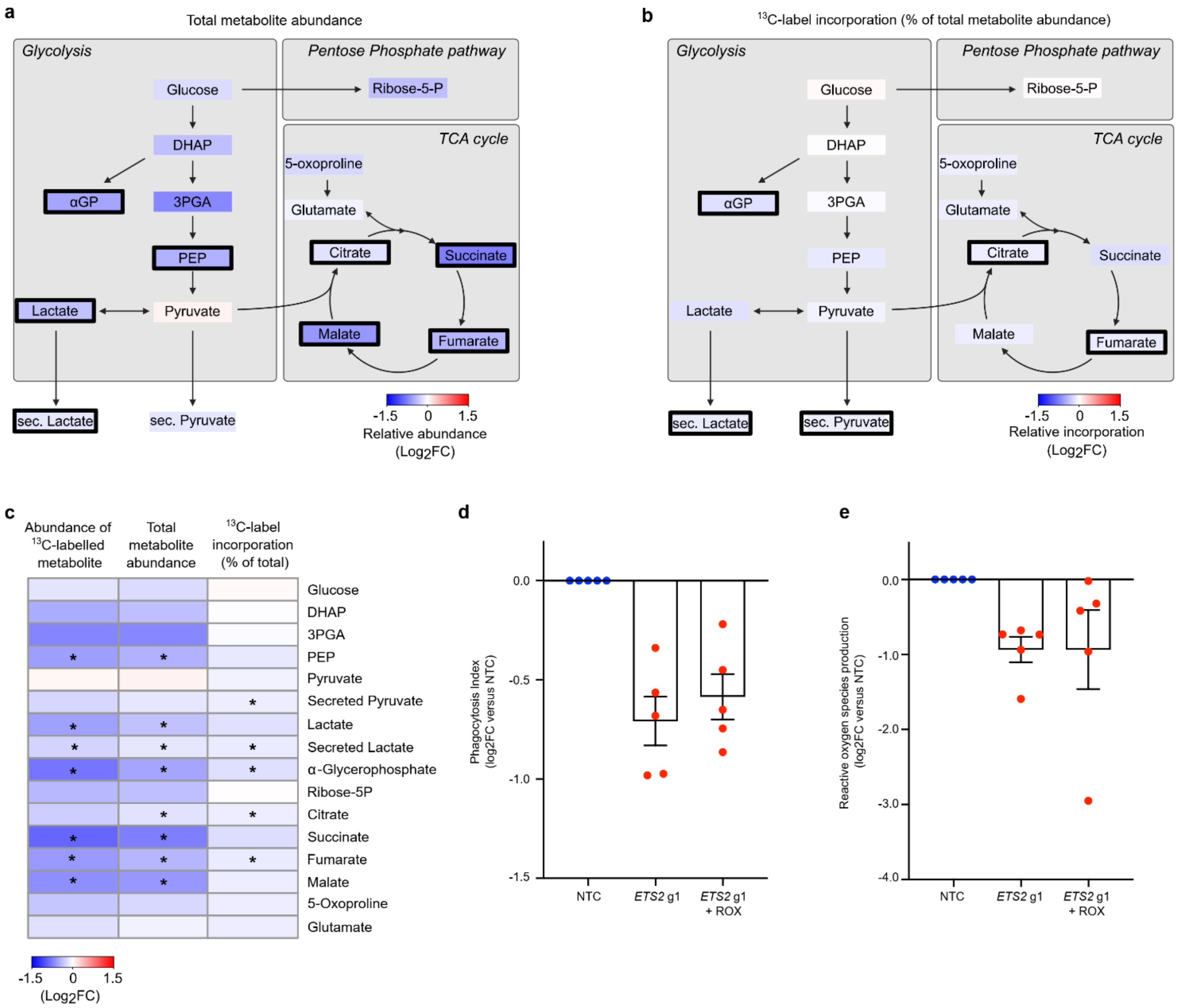


**Extended Data Figure 6. Metabolic effects of *ETS2* disruption.**

**a and b.** Changes in total metabolite abundance (**a**) and percentage of label incorporation from $^{13}$C-glucose (**b**) following *ETS2* editing in TPP macrophages (n=6). Colour depicts median log2 fold-change in *ETS2*-edited macrophages relative to unedited macrophages (transfected with non-targeting control RNPs; NTC). Bold black border indicates $P < 0.05$ (Wilcoxon signed rank test, two-tailed). **c.** Heatmap summarising metabolic changes following *ETS2* disruption. Colour depicts median log2 fold-change in *ETS2* g1-edited cells relative to unedited cells (Wilcoxon signed rank test, two-tailed). **d.** Phagocytosis index in unedited (NTC) and *ETS2*-edited TPP macrophages treated with roxadustat or vehicle. Phagocytosis index is calculated as proportion of positive cells multiplied by mean fluorescence intensity of positive cells (488 nm channel). Data normalised to phagocytosis index in unedited cells. **e.** Extracellular ROS production by unedited (NTC) and *ETS2*-edited TPP macrophages treated with roxadustat or vehicle – quantified using a chemiluminescence assay. Data represent log2 fold-change of area under curve (AUC) normalised to unedited (NTC) TPP macrophages. Error bars represent mean±SEM. * $P <$ 0.05.

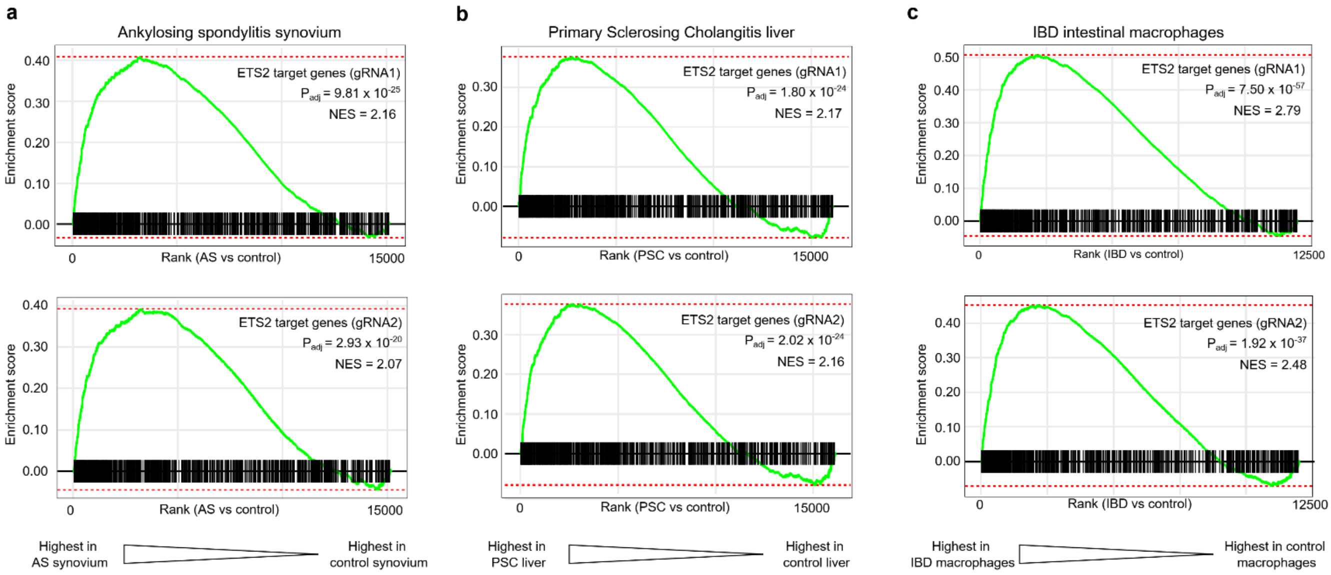


**Extended Data Figure 7. The transcriptional signature of *ETS2* is detectable in affected tissues from chr21q22-linked diseases.**

**a**. Gene set enrichment analysis (fGSEA) of ETS2-regulated genes within ankylosing spondylitis synovium (compared to control synovium). **b.** fGSEA of ETS2-regulated genes within primary sclerosing cholangitis liver biopsies (compared to control liver biopsies). **c**. fGSEA of ETS2-regulated genes within intestinal macrophages isolated from patients with active inflammatory bowel disease (compared to healthy control intestinal macrophages). ETS2-regulated gene sets in **a, b,** and **c** represent genes significantly downregulated following *ETS2* editing with gRNA1-containing RNPs (upper panel) or gRNA2-containing RNPs (lower panel).

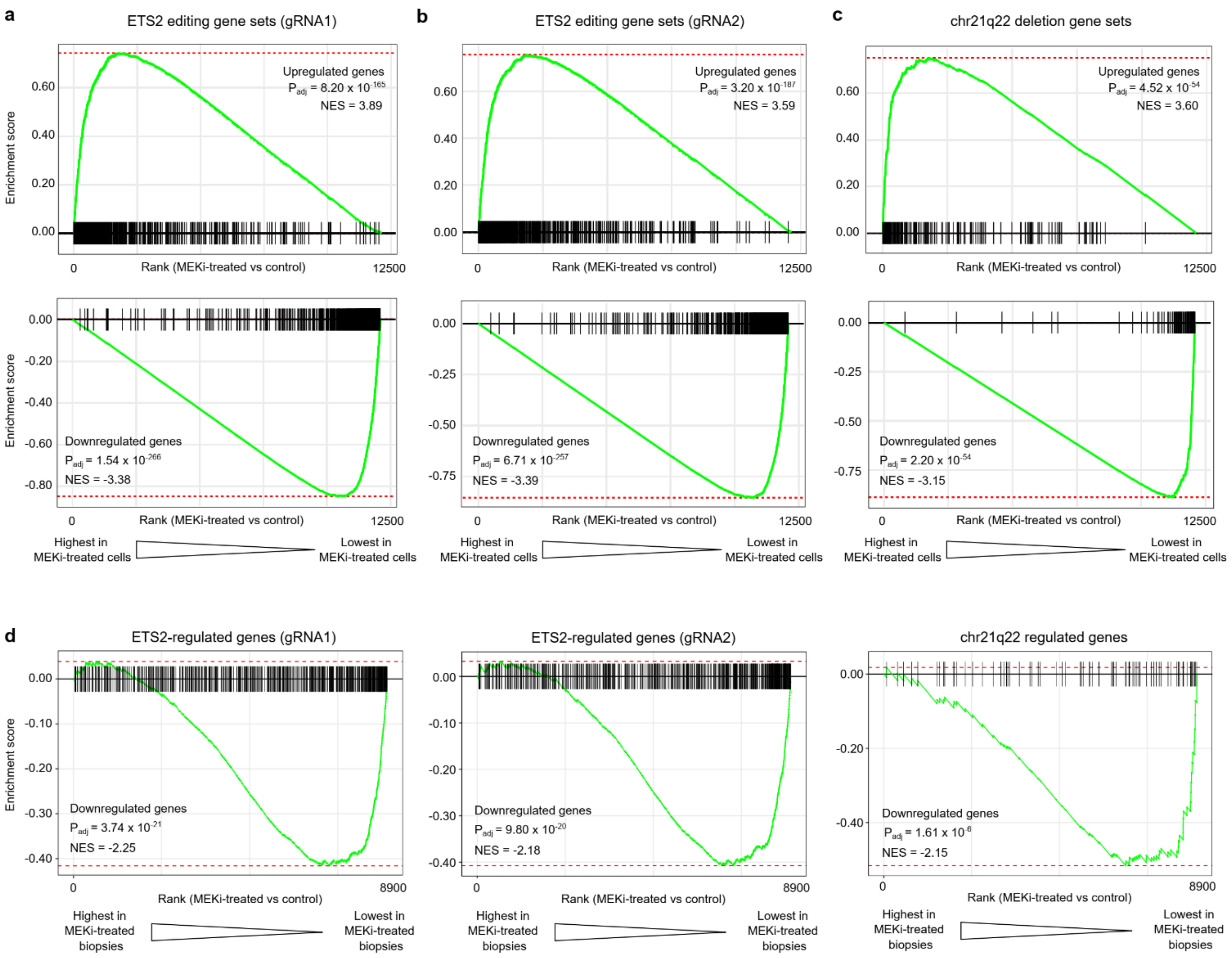


**Extended Data Figure 8. Effect of MEK1/2 inhibition on *ETS2*-regulated genes.**

**a, b and c**. Gene set enrichment analysis (fGSEA) in MEK1/2 inhibitor-treated TPP macrophages showing enrichment of gene sets upregulated (upper panel) or downregulated (lower panel) following *ETS2* or chr21q22 editing (MEK1/2 inhibited using PD-0325901, 0.5µM). Gene sets obtained from differential gene expression analysis (limma using voom transformation) following ETS2 disruption with gRNA1 (**a**), gRNA2 (**b**), or following chr21q22 deletion (**c**). **d.** fGSEA in intestinal biopsies from IBD patients showing enrichment of gene sets downregulated following *ETS2* or chr21q22 editing in MEK inhibitor-treated biopsies. Upregulated gene sets were not enriched.

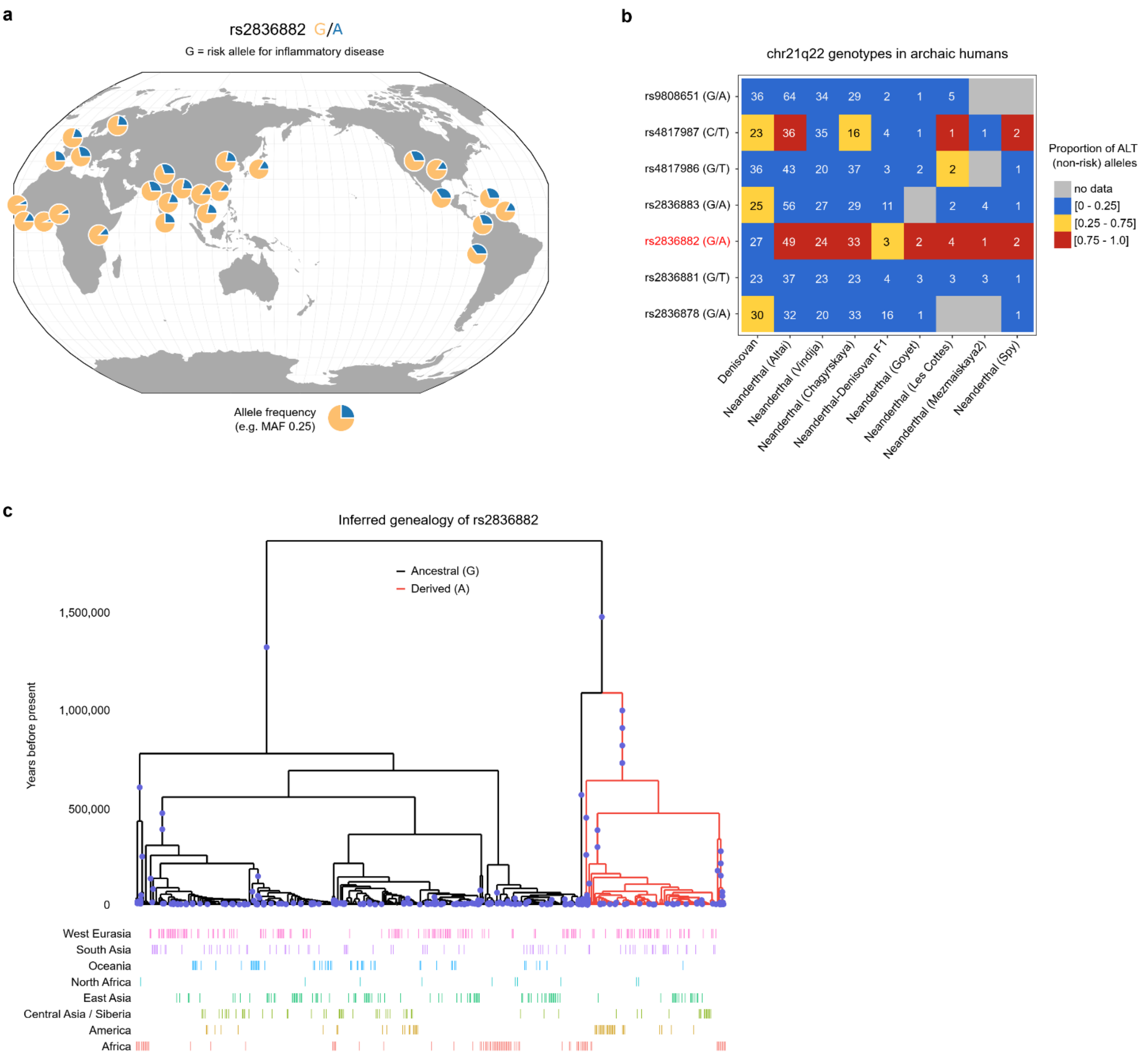


**Extended Data Figure 9. Geographic distribution and history of rs2836882.**

**a.** rs2836882 allele frequency in modern global populations (data from 1000 Genomes Project, plotted using Geography of Genetic Variants browser: https://popgen.uchicago.edu/ggv/). **b.** Genotypes of candidate SNPs at chr21q22 (99% credible set) in archaic humans (Neanderthals and Denisovans). Colour depicts the proportion of reads containing ALT alleles, with a value close to 0 consistent with a homozygous REF (risk) genotype, a value close to 1 consistent with a homozygous ALT (non-risk) genotype, and an intermediate value indicating a potential heterozygous genotype. Number in each cell indicates the number of reads at that SNP in the indicated sample. Putative causal variant highlighted in red. **c.** Inferred genealogy of the age of the rs2836882 polymorphism – analysed using Relate.

| | | | ETS2 gRNA 1 | | | ETS2 gRNA 2 | | |
|---|---|---|---|---|---|---|---|---|
| **Ensembl ID** | **Gene ID** | **PMID** | **logFC** | ***P*** | **adj.P.Val** | **logFC** | ***P*** | **adj.P.Val** |
| ENSG00000164691 | *TAGAP* | 23128233 | -0.91 | 7.55E-08 | 8.48E-05 | -1.00 | 1.53E-08 | 6.18E-05 |
| ENSG00000005844 | ***ITGAL*** | 28658209 | -0.77 | 3.32E-07 | 1.29E-04 | -0.72 | 8.99E-07 | 5.09E-04 |
| ENSG00000179630 | ***LACC1*** | 23128233 | -0.73 | 4.04E-07 | 1.29E-04 | -0.65 | 2.18E-06 | 6.71E-04 |
| ENSG00000163735 | *CXCL5* | 23128233 | -1.14 | 1.10E-06 | 2.78E-04 | -0.90 | 3.17E-05 | 2.29E-03 |
| ENSG00000172575 | ***RASGRP1*** | 23128233 | -0.70 | 5.85E-05 | 2.88E-03 | -0.70 | 5.90E-05 | 3.10E-03 |
| ENSG00000197943 | ***PLCG2*** | 36038634 | -0.25 | 1.49E-03 | 1.94E-02 | -0.33 | 7.02E-05 | 3.51E-03 |
| ENSG00000081237 | ***PTPRC*** | 26192919 | -0.54 | 7.11E-05 | 3.32E-03 | -0.51 | 1.52E-04 | 5.46E-03 |
| ENSG00000158714 | ***SLAMF8*** | 36038634 | -0.56 | 1.16E-04 | 4.53E-03 | -0.54 | 1.80E-04 | 5.86E-03 |
| ENSG00000108691 | *CCL2* | 23128233 | -0.83 | 1.21E-04 | 4.60E-03 | -0.79 | 2.10E-04 | 6.51E-03 |
| ENSG00000163110 | ***PDLIM5*** | 36038634 | -0.24 | 1.77E-03 | 2.17E-02 | -0.29 | 2.22E-04 | 6.71E-03 |
| ENSG00000136869 | ***TLR4*** | 26974007 | -0.49 | 6.73E-04 | 1.20E-02 | -0.54 | 2.22E-04 | 6.71E-03 |
| ENSG00000148400 | *NOTCH1* | 23128233 | -0.44 | 1.41E-03 | 1.88E-02 | -0.52 | 2.72E-04 | 7.50E-03 |
| ENSG00000134242 | ***PTPN22*** | 28658209 | -0.40 | 2.66E-03 | 2.77E-02 | -0.48 | 5.70E-04 | 1.16E-02 |
| ENSG00000079263 | ***SP140*** | 26192919 | -0.42 | 6.09E-03 | 4.72E-02 | -0.52 | 9.96E-04 | 1.64E-02 |
| ENSG00000169403 | ***PTAFR*** | 36038634 | -0.71 | 4.50E-04 | 9.70E-03 | -0.65 | 1.08E-03 | 1.71E-02 |
| ENSG00000150637 | *CD226* | 23128233 | -0.43 | 2.05E-03 | 2.36E-02 | -0.46 | 1.20E-03 | 1.82E-02 |
| ENSG00000143226 | ***FCGR2A*** | 23128233 | -0.99 | 2.01E-04 | 6.18E-03 | -0.79 | 1.93E-03 | 2.38E-02 |
| ENSG00000138821 | *SLC39A8* | 26192919 | -0.71 | 3.90E-04 | 8.98E-03 | -0.57 | 2.74E-03 | 2.93E-02 |
| ENSG00000100365 | ***NCF4*** | 36038634 | -0.59 | 5.26E-04 | 1.05E-02 | -0.47 | 3.77E-03 | 3.54E-02 |
| ENSG00000187796 | ***CARD9*** | 23128233 | -0.51 | 1.18E-03 | 1.67E-02 | -0.44 | 3.89E-03 | 3.61E-02 |
| ENSG00000121281 | ***ADCY7*** | 28067910 | -0.32 | 3.67E-03 | 3.42E-02 | -0.32 | 3.96E-03 | 3.66E-02 |
| ENSG00000144802 | ***NFKBIZ*** * | 26192919 | -0.28 | 1.55E-03 | 2.00E-02 | -0.23 | 7.10E-03 | 5.26E-02 |
| ENSG00000167207 | ***NOD2*** * | 28658209 | -0.78 | 3.94E-04 | 9.01E-03 | -0.53 | 9.66E-03 | 6.32E-02 |

**Extended Data Table 1. IBD risk genes downregulated following *ETS2* disruption.**

Results shown for IBD-associated genes that were differentially expressed between *ETS2*-edited and unedited (NTC) TPP macrophages (n=9).
Log2 fold-change is with respect to expression in unedited cells.
Genes in bold have been denoted as causal at their respective loci or are the only candidate gene at the locus.
*P*-value adjusted for multiple testing using Benjamini-Hochberg method.
PMID denotes PubMed ID of study with strongest IBD association for each gene.
* consistent effect but adjusted *P* value (adj.P.val) < 0.05 for one gRNA only.
